# Direct non-transcriptional link between brassinosteroid perception and cortical microtubule reorientation drives hypocotyl growth

**DOI:** 10.1101/2024.07.12.603229

**Authors:** Charlotte Delesalle, Alvaro Montiel-Jorda, Julie Neveu, Satoshi Fujita, Grégory Vert

**Affiliations:** Plant Science Research Laboratory (LRSV), UMR5546 CNRS/University of Toulouse/Toulouse-INP, 24 chemin de Borde Rouge, 31320 Auzeville Tolosane, France

## Abstract

Brassinosteroids (BRs) are steroid-type phytohormones that are essential for plant growth, development and adaptation to environmental stresses. BRs promote cell wall remodelling through extensive reprogramming of gene expression, thus impacting on cell elongation and hypocotyl growth. The existence of non-genomic responses to BRs is however still unclear. Here we identified and characterized a new family of BRI1-interacting proteins identified by yeast two hybrid named MBAPs. We confirmed using several complementary approaches that MBAPs are genuine BRI1 partners in plant cells. We demonstrated that MBAPs are localized to cortical microtubules (CMTs) underlying the plasma membrane using a short conserved helix and where they contact with and are phosphorylated by BRI1. Combinations of *mbap* loss-of-function mutants showed hypersensitivity to BRs and establish MBAPs as negative regulators of BR responses. Surprisingly, *mbap* mutants showed unaffected typical downstream BR signaling readouts and BR target gene expression. Rather, *mbap* mutants displayed disordered CMT network and enhanced CMT reorganization upon BR perception. Altogether, our work shed light on the direct non-transcriptional connection between BR perception at the cell surface and CMT organization in the control of hypocotyl elongation.

## Introduction

Brassinosteroids (BRs) are plant steroid hormones essential for many aspects of plant growth and development such as cell division, cell elongation, vascular tissue differentiation, reproduction processes, and stress responses (Fujioka et al., 1997; Li and Chory, 1997; Manghwar et al., 2022; Nolan et al., 2020; Wang et al., 2001). In particular, BRs are crucial for the growth of several plant organs including the embryonic stem called hypocotyl. Mutants impaired in BR biosynthesis or signaling show severe dwarfism, pointing to the BR growth-promoting effect. The BR signaling cascade and genomic responses are among the best characterized and understood pathway in plants. BRs are sensed at the plasma membrane (PM) by the BRASSINOSTEROID INSENSITIVE 1 (BRI1) family of Receptor-Like-Kinases (Clouse et al., 1996; Li and Chory, 1997), initiating signal transduction events that inhibits the BR-INSENSITIVE 2 (BIN2) kinase which acts as negative regulator of the pathway (He et al., 2002; Li et al., 2001). This in turns allows the dephosphorylation of BRI1-EMS-SUPPRESSOR 1 (BES1) and BRASSINAZOLE RESISTANT 1 (BZR1) family transcription factors to control expression of thousands of target genes. Among BR-regulated genes are found many genes related to cell wall synthesis, extensibility and growth. These include genes encoding CELLULOSE SYNTHASES (CESAs) involved in cell wall (CW) cellulose fibril deposition, CW-modifying enzymes allowing cell wall loosening and turgor pressure-mediated irreversible deformation, and microtubules that are the directional determinants of CESAs.

Microtubules are hollow biopolymers that consist of α,β-tubulin heterodimers. These filaments structure show high dynamicity, enabling to reorganize their structures and organization quickly (Hashimoto, 2015). Interphase cortical microtubules (CMT) expand beneath the PMs and serve as guide rails of cellulose synthase complexes and target points for their integration into the PMs (Paredez et al., 2006; Crowell et al., 2009; Gutierrez et al., 2009;). As cellulose microfibrils provide a main load-bearing source for determining cell shapes, the control of CMT orientation is a critical step in plant cell growth (Cosgrove, 2005). CMT orientation is known to be controlled by internal or external physicochemical cues, such as light, physical force, salt, and phytohormones (Chen et al., 2016; Hamant et al., 2008; Lindeboom et al., 2013; Locascio et al., 2013; True and Shaw, 2020). CMT organization is often executed by MICROTUBULE-ASSOCIATED PROTEINS (MAPs) or direct modification of tubulins, depending on the types of triggers (Fujita et al., 2013; Lee et al., 2012; Lindeboom et al., 2018; Nakamura et al., 2018; Takatani et al., 2020).

The precise molecular mechanisms underlying the connections between BRs and microtubules are still largely unknown. BRs were shown to promote alignment of CMT to transverse axis through upregulating expression of gene encoding MICROTUBULE-DEPOLYMERIZING PROTEIN40 (MDP40) (Wang et al., 2012). The defects in CMT reorganization observed in *mdp40* loss-of-function mutants result in abnormal cell/organ growth. A few recent reports have however highlighted more direct links between BRs and CMTs. The MAP CLIP170-ASSOCIATED PROTEIN (CLASP) was shown to mediate BR-dependent CMT reorganization, facilitating BRI1 trafficking to the PM by tethering BRI1-containing vesicles to CMT through the interaction with SORTING-NEXIN1 (SNX1) (Ambrose et al., 2013; Ruan et al., 2018). BRs in turn negatively regulate *CLASP* transcription, suggesting that BR signaling and CMT reorganization by CLASP constitute a negative feedback loop. Besides the transcriptional regulations depicted above, BRs also impact on CMTs through non-genomic responses. ROP signaling indeed connects the BR signaling pathway and CMT organization since BIN2 phosphorylates and controls the activity of the PHGAP1/2 (PLECKSTRIN HOMOLOGY GTPase ACTIVATING PROTEINS) modulators of ROP signaling (Lauster et al., 2022; Zhang et al., 2022). This signaling ensures indent formation by locally stabilizing CMTs. Altogether, a growing amount of evidence points toward a close link between CMTs and BR signaling (Delesalle et al., 2024) but the underlying mechanisms driving such direct crosstalk are still largely unknown.

In this study, we uncovered a new family of MAPs that directly interact with BRI1, named MICROTUBULE-BRI1 ASSOCIATED PROTEINS (MBAPs). Genetic and pharmacological approaches revealed that MBAPs are novel negative regulators of BR-mediated hypocotyl elongation acting independently from the canonical downstream BR signaling pathway and typical BR-regulated gene regulation. MBAPs act as stabilizer of the CMT network and limit BR-induced CMT reorientation. Taken together, our work unraveled a new branch of the BR signaling pathway that drives hypocotyl cell elongation through the reorganization of the CMT network via the BRI1-MBAP axis.

## Results

### Identification of a new family of BRI1-interacting proteins

To identify novel proteins regulating BRI1, we performed a yeast two-hybrid screen using the BRI1 cytoplasmic domain comprising the juxtamembrane domain, the kinase domain, and the C-terminal tail as bait. The library used was prepared using RNA extracted from seven-day-old light-grown Arabidopsis seedlings, condition where BRI1 is highly expressed. Eighty-four million yeast transformants were screened for growth in selective media. This screen recaptured well-known BRI1 interactors such MEMBRANE-ASSOCIATED KINASE REGULATOR 1 (MAKR1), SOMATIC EMBRYOGENESIS RECEPTOR-LIKE KINASE 1 (SERK1), or TRANSTHYRETIN-LIKE PROTEIN (TTL) (Nam and Li, 2004, 2002; Santiago et al., 2013; Wang and Chory, 2006). In addition, three novel plant-specific proteins of unknown function and belonging to the same family were identified as interacting with BRI1 kinase domain. These proteins are annotated in the TAIR database as i) LOW protein: protein phosphatase 1 regulatory subunit-like protein (AT1G17360), ii) COP1-interacting protein-like protein (AT1G72410) and iii) GPI-anchored adhesion-like protein (AT3G14172). Sequence analyses between the three proteins revealed about 64 to 70% percent of identity over the whole protein sequence, and highlighted conserved motifs in N-terminus (N-Ter) with COP1-INTERACTING PROTEIN 7 (CIP7) (Arico et al., 2024; Yamamoto et al., 1998) and related proteins from *Medicago truncatula*, *Marchantia polymorpha* and *Oryza sativa* (Fig. 1A). Interestingly, the rice mutant for the ortholog *Os07g0616000* gene has been previously characterized and shows defects in seed or leaf development (Abe et al., 2010; Li et al., 2010; Zhu et al., 2010, Zhu et al., 2021). Due to the molecular features of these proteins that will be presented hereafter, we named these family members *MBAP1/2/3* for *MICROTUBULE-BRI1 ASSOCIATED PROTEINS*.

**Fig. 1.**
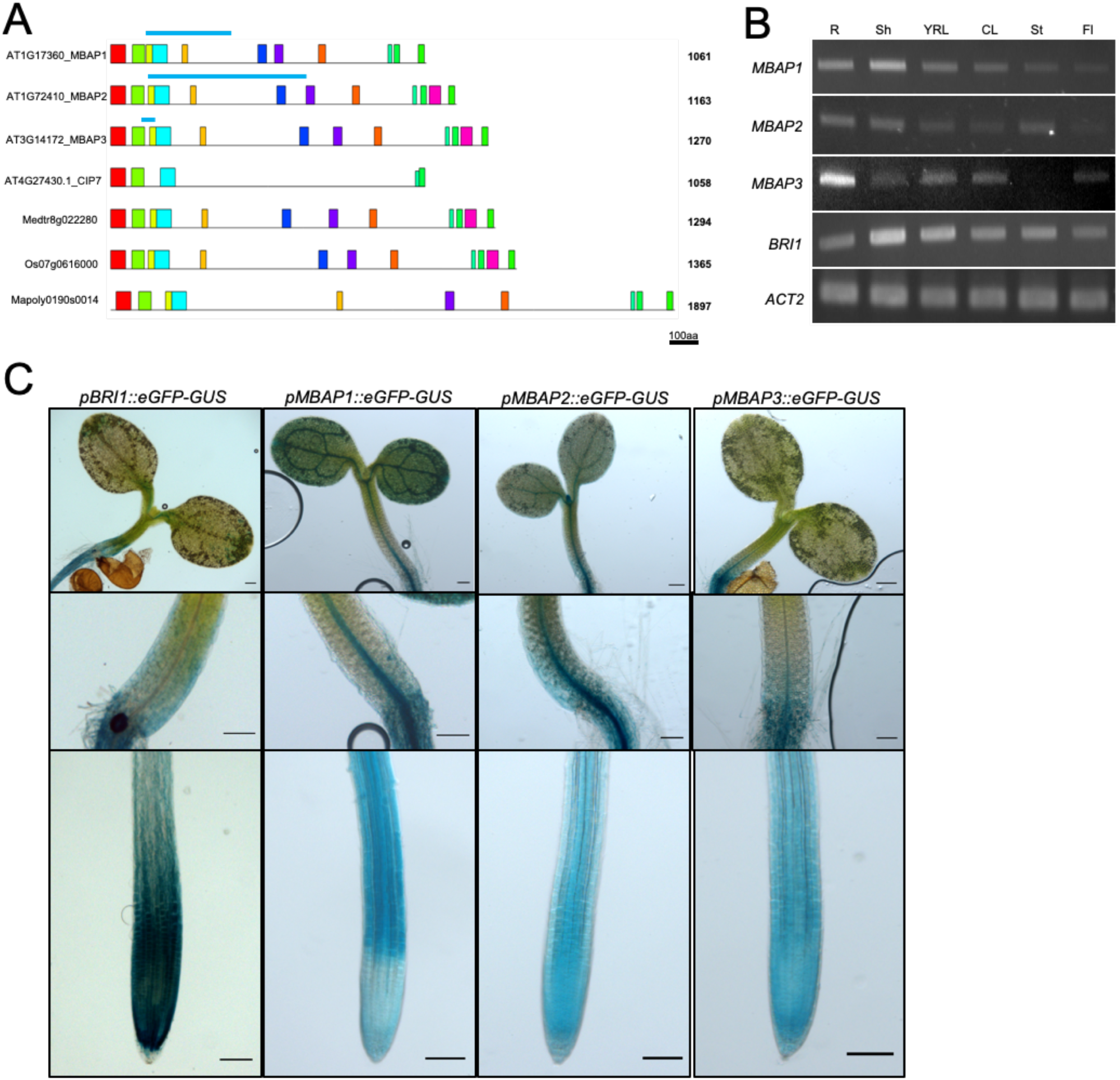
Identification of the MBAP family of proteins and expression pattern in Arabidopsis plants. (A) Schematic representation of the Arabidopsis MBAP1/2/3 protein sequences with their paralog *CIP7*, and their *M. truncatula*, *O. sativa*, *M. Polymorpha* orthologs. Colored boxes represent conserved motifs. Black lines indicate non-conserved regions. Regions of interaction with BRI1 identified by Y2H are depicted in blue, and reveal a short overlapping region common to MBAP1-3. (B) *MBAP1*, *MBAP2*, *MBAP3* and *BRI1* expression by RT-PCR in roots (R), shoots (Sh), young rosette leaves (YL), cauline leaves (CL), stems (St), and flowers (Fl). *ACT2* expression is used as a control. (C) Promoter activity of *BRI1* and *MBAP* promoters driving GUS expression in 5-day-old seedlings.

We first evaluated the expression pattern of *MBAP* genes in Arabidopsis. Using RT-PCR, *MBAP1/2/3* were shown to be widely expressed in whole plants, clearly overlapping with *BRI1* expression (Fig. 1B). To further characterize the expression profile of *MBAP* genes, transgenic plants expressing ß-glucuronidase (GUS) under the control of *MBAP* promoters were generated and analyzed. GUS activity driven by the *MBAP1-3* promoters was detected in the whole seedling, especially in cotyledons, hypocotyls, and roots and clearly intersected with *BRI1* promoter activity (Fig. 1C). Such expression profile for *MBAPs* overall matches where BR signaling is active and where *BRI1* expression was previously observed (Friedrichsen et al., 2000), supporting the possibility of MBAPs interacting with BRI1 in plants.

### BR receptors and MBAPs interact at the plasma membrane and at intracellular filaments

To validate *in planta* the BRI1-MBAP1 interaction, we first carried out coimmunoprecipitation assays using transiently-expressed functional BRI1-mCitrine (BRI1-mCit) (Jaillais et al., 2010) and MBAP1-mCherry (MBAP1-mChe) fusions. Additionally, leaf discs were treated with mock or epibrassinolide (eBL) to assess the impact of BRs on the BRI1-MBAP1 interaction. We used anti-GFP antibodies to immunoprecipitate BRI1-mCit or mCit alone as negative control. Immunoprecipitates from BRI1-mCit showed a band migrating at the expected size of BRI1 mCitrine fusion protein (170 kDa ; Martin et al., 2015) (Fig. 2A). MBAP1-mCherry successfully coimmunoprecipitated with BRI1-mCit but not with mCit, indicating MBAP1 specifically interacts with BRI1. Besides, BR treatment had no significant effect on the levels of MBAP1 coimmunoprecipitated with BRI1 (Fig. 2A, 2B). To ascertain that the BR treatment was effective in our conditions, we confirmed that BKI1 dissociates from the PM under such conditions, as previously reported in Arabidopsis (Wang and Chory, 2006; Jaillais et al., 2011) (Supplemental Fig. 1).

**Fig 2.**
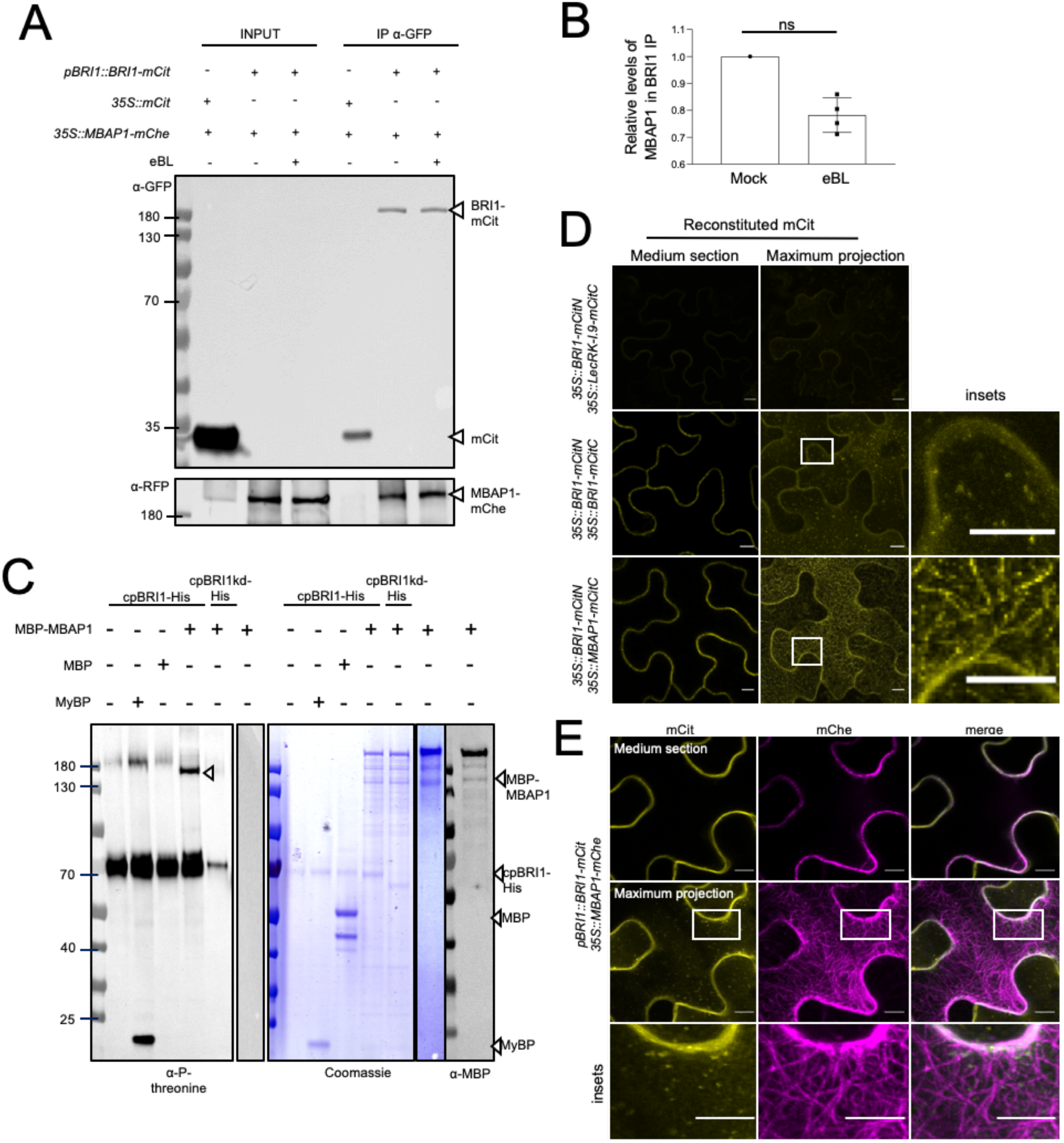
MBAP1/BRI1 interaction occurs at the PM and at filamentous structures. (A) Coimmunoprecipitation assay from *N. benthamiana* leaves coexpressing *pBRI1::BRI1-mCit* and *35S::MBAP1-mCherry* treated with 10µM eBL or mock for 20 min. *35S::mCitrine* was used as a negative control. (B) Quantification of relative coimmunoprecipitated MBAP1/BRI1 ratio in 4 replicates. Data are normalized to their mock counterpart (C) *In vitro* kinase assay using His-tagged cytoplasmic domain of BRI1 (cpBRI1-His) and MBP-tagged MBAP1 (MBP-MBAP1). Phosphorylated threonine residues were detected using anti-phospho-threonine antibodies. Coomassie staining is shown for loading control. His-tagged cytoplasmic domain of BRI1 kinase dead (cpBRI1kd-His), MBP alone and an artificial substrate, myelin basic protein (MyBP) were used as negative and positive controls, respectively. Immunoblots with anti-MBP antibodies shows the 3 bands corresponding to purified MBP-MBAP1. (D) Bimolecular Fluorescence Complementation (BiFC) in *N. benthamiana* coexpressing *35S::BRI1-mCitN* and *35S::MBAP1-mCitC,* or *35S::BRI1-mCitN* and *35S::LecRK-I.9-mCitC* as negative control. Coexpression of *35S::BRI1-mCitN and 35S::BRI1-mCitC* serves as positive control. Insets show higher magnification. (E) Colocalization of BRI1 and MBAP1 at the PM and in filamental structures in leaf discs of *N. benthamiana* expressing *pBRI1::BRI1-mCitrine* and *35S::MBAP1-mCherry*. The upper half of cells is shown as maximum projection of Z-stacks. Scale bar: 10 µm. Insets show higher magnification.

To further validate the association of MBAP1 and BRI1 and its functional relevance, we tested if MBAP1 was a substrate of BRI1 kinase activity. To this purpose, we performed *in vitro* kinase assay using bacterially-expressed C-terminally His tag-fused BRI1 cytoplasmic domain (cpBRI1-His), or its kinase-dead version (cpBRI1kd-His) as negative control, and MBAP1 fused to the maltose-binding protein (MBP-MBAP1). A strong auto-phosphorylation of BRI1 was detected in cpBRI1-His alone using anti-phospho-threonine antibodies (P-Thr), while no signal was observed for the kinase-dead cpBRI1kd-His (Fig. 2C). Coincubation of cpBRI1-His with MyBP (Myelin basic proteins), a widely-used generic kinase substrate, confirmed the trans-phosphorylation activity of BRI1 kinase. Additionally, cpBRI1-His was unable to transphosphorylate MBP alone, while a phosphorylation signal was observed for MBP-MBAP1 (expected size 162 kDa). Of note, purification of MBP-MBAP1 always resulted in 3 distinct bands with the lower band corresponding to the expected size of MBP-MBAP1. Again, no signal was observed for MBP-MBAP1 when incubated with cpBRI1kd. Altogether, these results indicate that BRI1 specifically phosphorylates MBAP1 *in vitro*.

To visualize the BRI1-MBAP1 interaction, we then performed bimolecular functional complementation (BiFC) assay using transiently expressed fusion proteins of BRI1 and MBAP1 with the N- and C-terminal fragments of mCit (mCitN and mCitC), respectively. We used the reported BRI1 homodimerization as positive control (Wang et al., 2005), and the lectin-RLK LecRK-I.9 (also known as DORN1) as negative control (Bouwmeester et al., 2011; Choi et al., 2014). As expected, no fluorescence was observed when BRI1-mCitN was coexpressed with LecRK-I.9-mCitC while coexpression of BRI1-mCitN and BRI-mCitC resulted in strong signal at the PM and in dots likely corresponding to endosomes. mCit fluorescence resulting from the BRI1-MBAP1 interaction was found at the PM when the median focal plane was imaged (Fig. 2D). Consistently, BRI1 and MBAP1 proteins both colocalized at the PM when (Fig. 2E). Interestingly, the BRI1-MBAP1 interaction visualized by BiFC using maximum projection from the whole cell also occurred at small intracellular filaments (Fig. 2D), where MBAP1-mCherry is also found, and possibly colocalizes with BRI1-positive endosomes (Fig. 2E). A similar BiFC pattern was observed for the BRI1-MBAP2 or BRI1-MBAP3, supporting the fact that all three MBAP proteins likely interact with BRI1 in plants (Supplemental Fig. 2). In addition, the BRI1 homologs BRL1 and BRL3 that are also involved in BR perception also interacted with MBAP1-3 (Supplemental Fig. 2).

Taken together, our results indicate that MBAP1 is an *in vitro* substrate of BRI1 and that MBAP proteins interact with BR receptors at the cell surface and possibly on intracellular filaments.

### MBAPs are microtubule-associated proteins

In order to unveil the nature of the intracellular filaments where MBAP1 is observed and BRI1-MBAP1 interaction occurs, we colocalized transiently-expressed MBAP1 with intracellular markers. When coexpressed with the Microtubule Binding Domain of MAP4 GFP-MBD_MAP4_ MT marker (Ruan et al., 2018) in *N. benthamiana* leaves, MBAP1-mChe fluorescence clearly overlapped with GFP-MBD_MAP4_ (Fig. 3A), indicating that MBAP1 is found associated to MTs. MBAP2 and MBAP3 were also found to colocalize with the MBD_MAP4_ (Supplemental Fig. 3A). The presence of MBAP1 to MTs was further confirmed by the destabilization of MBAP1 localization upon treatment with the oryzalin MT network depolymerizing drug (Nakamura et al., 2004) (Fig. 3A). To validate such observations, we sought to generate stable Arabidopsis transgenic lines expressing GFP-tagged MBAP1 but corresponding transgenics rapidly cosuppressed, regardless of the promoter, tag or tag position used. We however managed to obtain evidence that MBAP1 colocalizes with the mScarlet-tagged TUBULIN6 (mSca-TUB6) in stable transgenic lines (Supplemental Fig. 3B).

**Fig 3.**
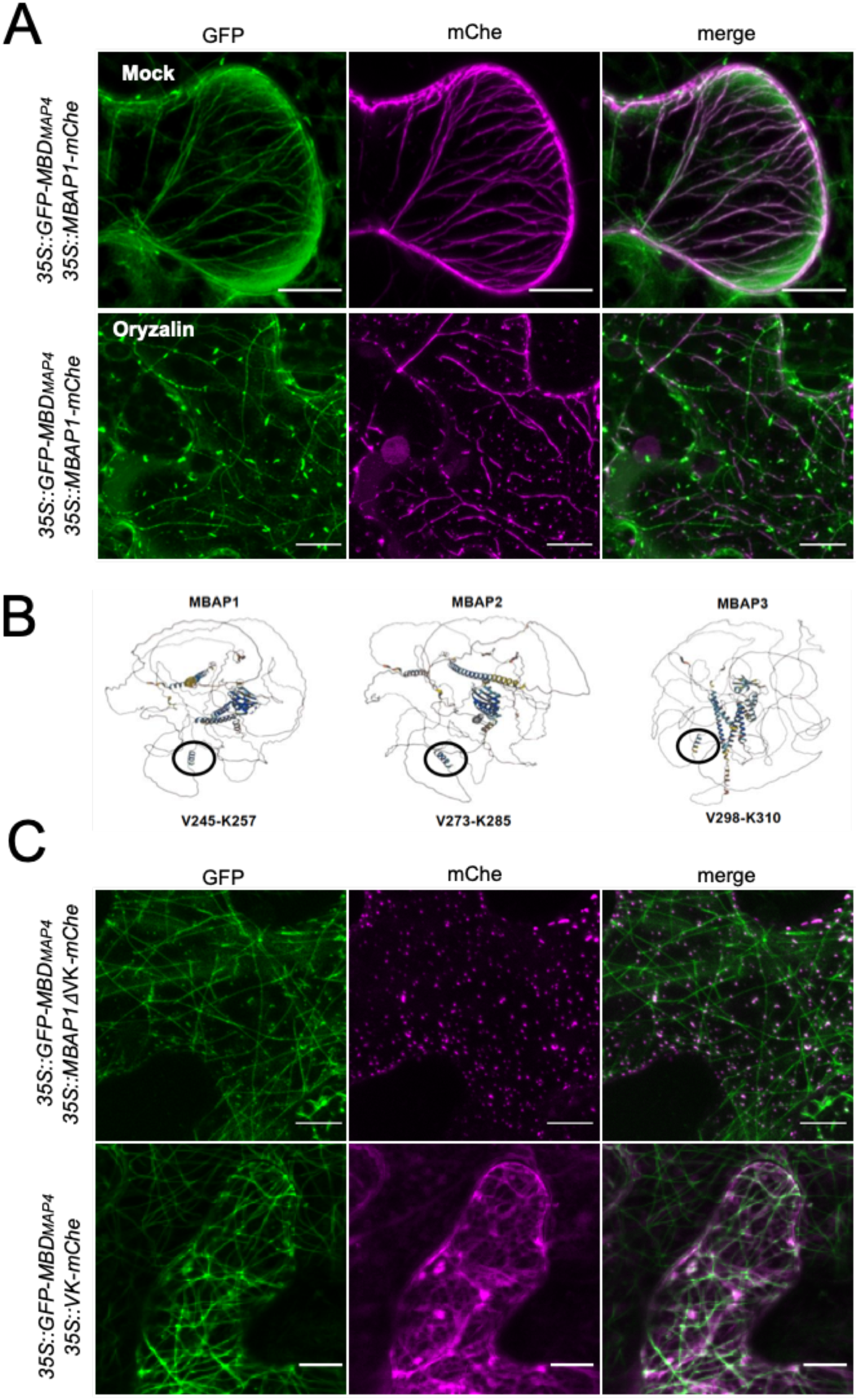
MBAP1 localizes to microtubules using a short helix. (A) MBAP1 colocalizes with the MBD_MAP4_ microtubule marker and is sensitive to oryzalin. *N. benthamiana* leaf discs coexpressing *35S::MBAP1-mCherry* and *p35S::GFP-MBDMAP4* treated with mock (top) or 10 µM oryzalin for 15 min (bottom) were observed by confocal microscopy. Representative pictures are shown as maximum projection of Z-stacks. Scale bar: 10 µm. (B) Alphafold protein structure prediction for MBAP1, MBAP2 and MBAP3. Circles highlight a highly conserved helix of 12 aa in MBAP proteins. Position of the helix is specified for each MBAP protein. (C) The VK helix is necessary for localization of MBAP1 to MTs. *N. benthamiana* leaf discs coexpressing 35S::GFP-MBD_MAP4_ and 35S::MBAP1ΔVK-mCherry were observed by confocal microscopy. Panels correspond to maximum projection of Z-stacks. Scale bar: 10 µm. (D) The VK helix is sufficient for localization to MTs. *N. benthamiana* leaf discs coexpressing 35S::GFP-MBD_MAP4_ and 35S::VK-mCherry were observed by confocal microscopy. Panels correspond to maximum projection of Z-stacks. Scale bar: 10 µm.

We next addressed whether the association between MTs and MBAP1 is direct or indirect using an *in vitro* MT-centrifugation assay. The full-length version of MBP-MBAP1 precipitated without MTs, thus preventing us from using it in MT centrifugation assays. We therefore decided to use fragment versions of MBAP1 that colocalize with the MT marker. We generated MBAP1 truncations (MBAP1_1-190_, MBAP1_191-489_ and MBAP1_490-1061_) and fused them to mCherry. Confocal microscopy analyses using transiently-expressed MBAP1-mChe truncations showed that only MBAP1_191-489_-mChe displayed the same filamentous pattern as observed for MBAP1 partially colocalizing with MTs (Supplemental Fig. 4A), suggesting that such fragment contains the MT-binding domain. We thus produced and purified MBP-MBAP1_191-489_ recombinant protein from *E coli* and performed the *in vitro* MT-centrifugation assay. When incubated with MTs, the negative (BSA) and positive (MAP fraction) controls were found in cytosol and pellet, respectively (Supplemental Fig. 4B). Regardless of the presence of MTs, MBP-MBAP1_191-489_ was recovered in the supernatant, suggesting that MBAP1 association with MTs is likely not direct or that direct binding to MTs requires additional features occurring in live cells such as dimerization or post-translational modifications.

Because MT association domain is likely due to secondary structures, as shown for MAP65-1 (Li et al., 2007) we used the Alphafold structure prediction of MBAPs to identify secondary structures involved in MT recruitment. MBAPs are highly disordered proteins but show a structured region in their N-terminus. Of particular interest is a small helix of 12 amino acids located between residues V245 and K257 and found in the MBAP1_191-489_ fragment and that is shared with MBAP2 and MBAP3 (Fig. 3B). To examine the functional role of such helix, we first generated an MBAP1 variant called MBAP1ΔVK lacking the corresponding amino acids. Transient expression of MBAP1ΔVK-mChe with MBD_MAP4_-GFP revealed an absence of MT localization for MBAP1ΔVK (Fig. 3C), with only cytosolic dots being observed. This indicates that this short helix is necessary for MT recruitment of MBAP1. To evaluate if the VK helix is also sufficient for MT recruitment, we investigated the localization of the sole VK helix fused to mChe and coexpressed it with MBD_MAP4_-GFP. VK-mChe displayed a filamentous pattern that overlapped, at least in part, with the MT marker (Fig. 3C). This argues for the VK helix being both necessary and sufficient for MT targeting of MBAP1.

Overall, our results clearly demonstrate that MBAPs is a new family of MAPs in *A. thaliana* using a short conserved helix for indirect recruitment to MTs.

### MBAPs inhibit the BR-mediated hypocotyl growth independently from the canonical BR signaling pathway

To investigate the biological role of MBAP family members and their functional relevance to BR signaling, we isolated two T-DNA insertion lines for each *MBAP* gene (Supplemental Fig. 5A). RT-PCR analyses confirmed that all T-DNA alleles did not express corresponding full-length transcripts (Supplemental Fig. 5B). No detectable macroscopic phenotype under normal growth conditions were observed in single *mbap* mutants. We next evaluated the sensitivity of light-grown *mbap* mutants to exogenously applied eBL. eBL treatment promotes hypocotyl elongation of light-grown Arabidopsis seedlings (Fig. 1A ; Supplemental Fig. 6). Single *mbap* mutants displayed wild-type sensitivity to BRs (Supplemental Fig. 6), suggesting that MBAPs have redundant functions. We therefore generated the 3 combinations of double mutants by crossing single T-DNA mutants, and confirmed absence of corresponding transcripts (Supplemental Fig. 5B). Compared to wild-type plants, all double mutant combinations showed hypersensitivity to BRs with longer hypocotyls on media supplemented with eBL, and resistance to the BR synthesis inhibitor propiconazole (PPZ) (Hartwig et al., 2012) (Fig. 4A). The *mbap2-mbap3* combination showed the most BR hypersensitive phenotype. To confirm that the lack of MBAP caused the BR-related phenotypes of *mbap* double mutant combinations, we expressed the best characterized MBAP1 member as a GFP-MBAP1 fusion in *mbap2-mbap3* and evaluated the BR responses of the corresponding plants. The *mbap2-mbap3/GFP-MBAP1* displayed wild-type responses to eBL (Fig. 4B), demonstrating that the loss of MBAP proteins yields BR hypersensitivity.

**Fig. 4.**
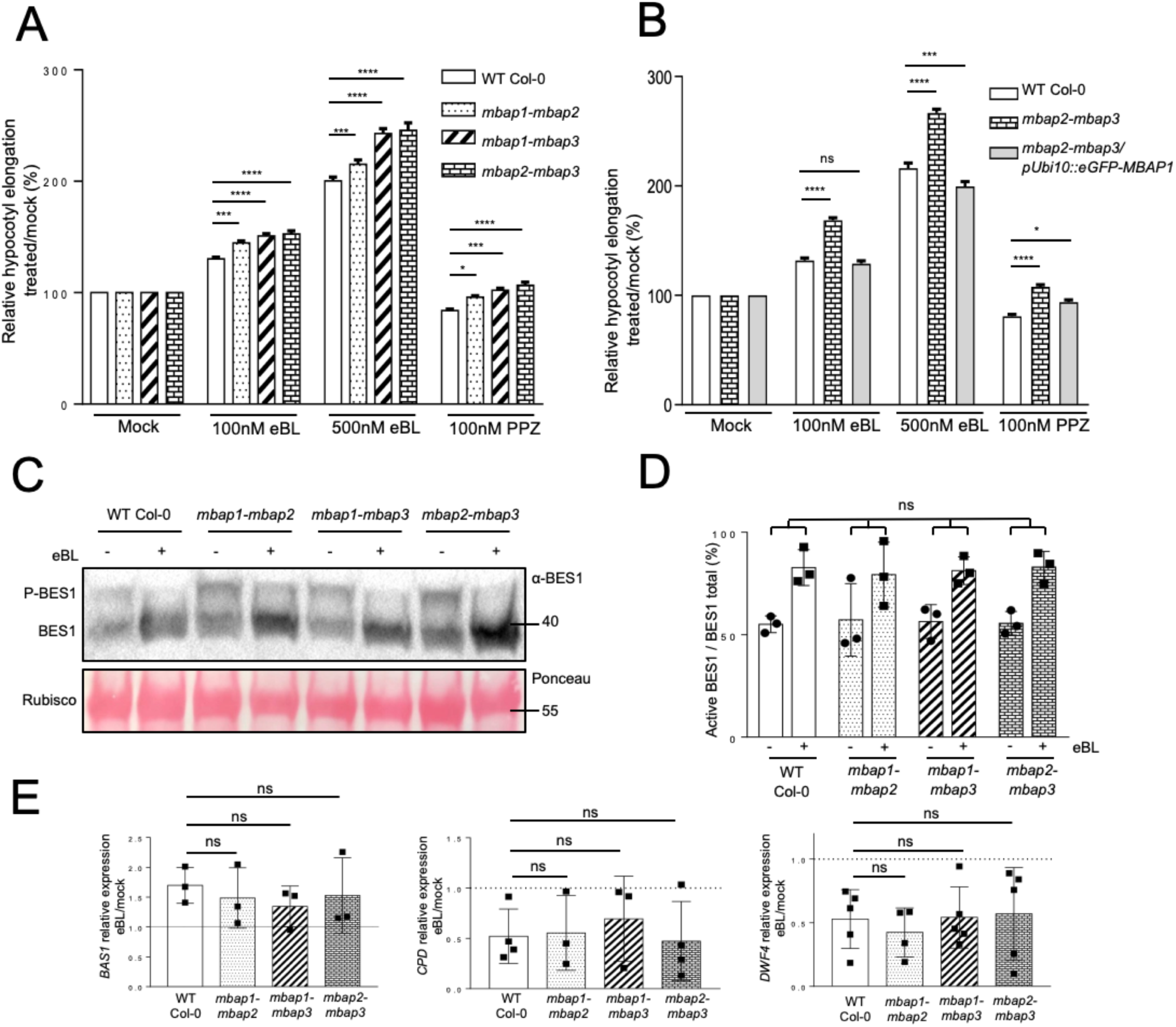
MBAP2/3 negatively regulate BR-induced hypocotyl elongation independently of the canonical BR pathway. (A) Relative hypocotyl length of 7-day-old WT, *mbap1-mbap2*, *mbap1-mbap3* and *mbap2-mbap3* seedlings on half-strength LS media containing epibrassinolide (100 nM, 500 nM eBL) or propiconazole (100 nM PPZ) compared to mock. Error bars represent SEM (n>30, 3 replicates). Asterisks represent statistically significant differences (p-value < 0.05 (*), 0.01 (**), 0.001 (***), 0.0001 (****) ; Two-way ANOVA with Dunnett’s multiple comparison). (B) Relative hypocotyl length of 7-day-old seedlings WT, *mbap1-mbap3*, *mbap2-mbap3*, and complemented *mbap2-mbap3/pUbi10::eGFP-MBAP1* on half-strength LS media containing epibrassinolide (100 nM, 500 nM eBL) or propiconazole (100 nM PPZ) compared to mock. Error bars represent SEM (n>30, 3 replicates). Asterisks represent statistically significant differences (p-value < 0.05 (*), 0.01 (**), 0.001 (***), 0.0001 (****) ; Two-way ANOVA with Dunnett’s multiple comparison). (C) Representative blot showing the phosphorylation status of BES1 in 7-day-old WT, *mbap1-mbap2*, *mbap1-mbap3,* and *mbap2-mbap3* treated with mock or 1µM eBL for 20 minutes. Phospho-BES1 (P-BES1; top) and dephospho-BES1 (BES ; bottom) were detected using anti-BES1 antibodies on total protein extract. Ponceau staining is used as loading control. (D) Quantification of the ratio between dephosphorylated BES1 and total BES1. Experiments were done in triplicates and error bars indicate SD (n=3). Mann-Whitney statistical test showed no significant difference between WT and mutants in all conditions. (E) Relative expression of BR-regulated genes *BAS1*, *CPD,* and *DWF4* analyzed by RT-qPCR after treating with mock or 1µM eBL for 3h. Data is shown as relative to mock-treated shown as the dotted line set to 1. Experiments were done in triplicates and error bars represent SD (n>3). Mann-Whitney statistical test showed no significant difference to the wild-type.

The BRI1-MBAP1 interaction combined to the BR hypersensitivity of double mutant *mbap* combinations suggest that loss of MBAPs yield enhanced BR signaling. We therefore analyzed typical molecular readouts of the BR signaling pathway such as the phosphorylation status of BES1 transcription factor (Yin et al., 2002) and expression changes of well-known BR-regulated genes (Vert et al., 2005). Exogenous BR treatment or plants hypersensitive to BRs trigger the accumulation of the fast-migrating dephosphorylated form of BES1 in immunoblot experiments using anti-BES1 antibodies (Yin et al., 2002). Wild-type and *mbap* mutant plants were therefore exposed to mock or eBL and subjected to immunoblot analyses using anti-BES1 antibodies. BRs promoted the dephosphorylation of BES1 in wild-type plants, as expected (Fig. 4C). Unexpectedly, *mbap* double mutants showed similar response to wild-type plants. Quantification of the ratio between dephosphorylated and phosphorylated forms of BES1 in 3 independent replicates confirmed that *mbap* double mutants showed a wild-type response to BRs when monitoring the activation of BES1 (Fig. 4D). To further examine the downstream BR responses of *mpab* mutants, we performed RT-qPCR analyses to monitor the expression of well-characterized BR-responsive genes. The BR biosynthetic enzyme genes *DWF4* and *CPD* are under negative feedback regulation by BRs, leading to reduced transcript accumulation upon BR exposure (Choe et al., 1998; Ohnishi et al., 2012). Conversely, the expression of the *BAS1* BR metabolic enzyme-encoding gene is induced following BR perception (Neff et al., 1999). Wild-type plants showed the typical expression profile for *CPD*, *DWF4* and *BAS1* when challenged with eBL (Fig. 4E). No statistical difference could be observed for *mbap* double mutant combinations compared to wild-type, further indicating that the phenotypic BR hypersensitivity caused by loss of *MBAP* genes is not caused by activation of the canonical BR signaling pathway and typical target gene expression (Vert et al., 2005).

Altogether, we demonstrate that MBAPs function genetically as negative regulators for BR-induced hypocotyl elongation independently of BR-triggered transcriptional regulation, pointing to the existence of extra BR-dependent routes contributing to hypocotyl growth.

### MBAPs stabilize the CMT network and inhibit BR-induced CMT reorientation

CMT organization and orientation is critical to establish the cell growth direction axis (Crowell et al., 2009; Gutierrez et al., 2009). BRs were previously reported to yield transversal reorientation of CMTs (Liu et al., 2018; Ruan et al., 2018; Wang et al., 2012) through transcriptional regulation of MAPs. Considering the localization of MBAP proteins to MTs and their ability to interact with BR receptors and convey BR responses, we hypothesized that BRs may act on CMTs through the MBAPs to regulate hypocotyl elongation. We first studied the impact of loss of MBAPs on the CMT network organization under resting conditions. To this purpose, the GFP-MBD_MAP4_ microtubule marker was crossed to the *mbap2-mbap3* double mutant. Confocal microscopy analyses of *mbap2-mbap3/GFP-MBD_MAP4_* revealed a marked difference in CMT organization (Fig. 5A). Quantification of CMT isotropy using Fibriltool plugin (Boudaoud et al., 2014) highlighted disordered and random CMT organization in *mbap2-mbap3* compared to wild-type (Fig. 5B), suggesting that MBAPs likely act as stabilizer of MTs or MT network. We then assessed the sensitivity of *mbap2-mbap*3/GFP-MBD_MAP4_ plants to oryzalin. The CMT network of WT plants was only mildly compromised after 20 min of oryzalin exposure (Fig. 5C, 5D). In sharp contrast, the combined loss of *MBAP2* and *MBAP3* led to a rapid and dramatic disappearance of the CMT network. These results confirmed that MBAPs are required to stabilize the CMT network in Arabidopsis hypocotyls. We next monitored the previously reported BR-mediated CMT reorientation in both wild-type and *mbap2-mbap3*. Confocal microscopy imaging and quantification of the mean filament angle by cell using Fibriltool showed that the BR mediated reorientation of CMTs in *mbap2-mbap3* is enhanced and faster than the one observed in wild-type plants, while no difference was detected in the absence of eBL (Fig. 5E, 5F). The increased reorientation of CMT network upon BRs is not associated with a change in MBAP1 localization since MBAP1-mCherry localization to CMTs appeared unaffected by BR treatment (Supplemental Fig. 7). CMT reorientation was long associated to cell elongation. We therefore measured cell expansion before and after BR treatment, yielding thus the rate of elongation in wild-type and *mbap2-mbap3*. Hypocotyl cell elongation rate upon exposure to BRs was significantly higher in *mbap2-mbap3* than in wild-type (Fig. 5G), consistent with the hypersensitivity to BRs we have observed (Fig. 4A).

**Fig 5.**
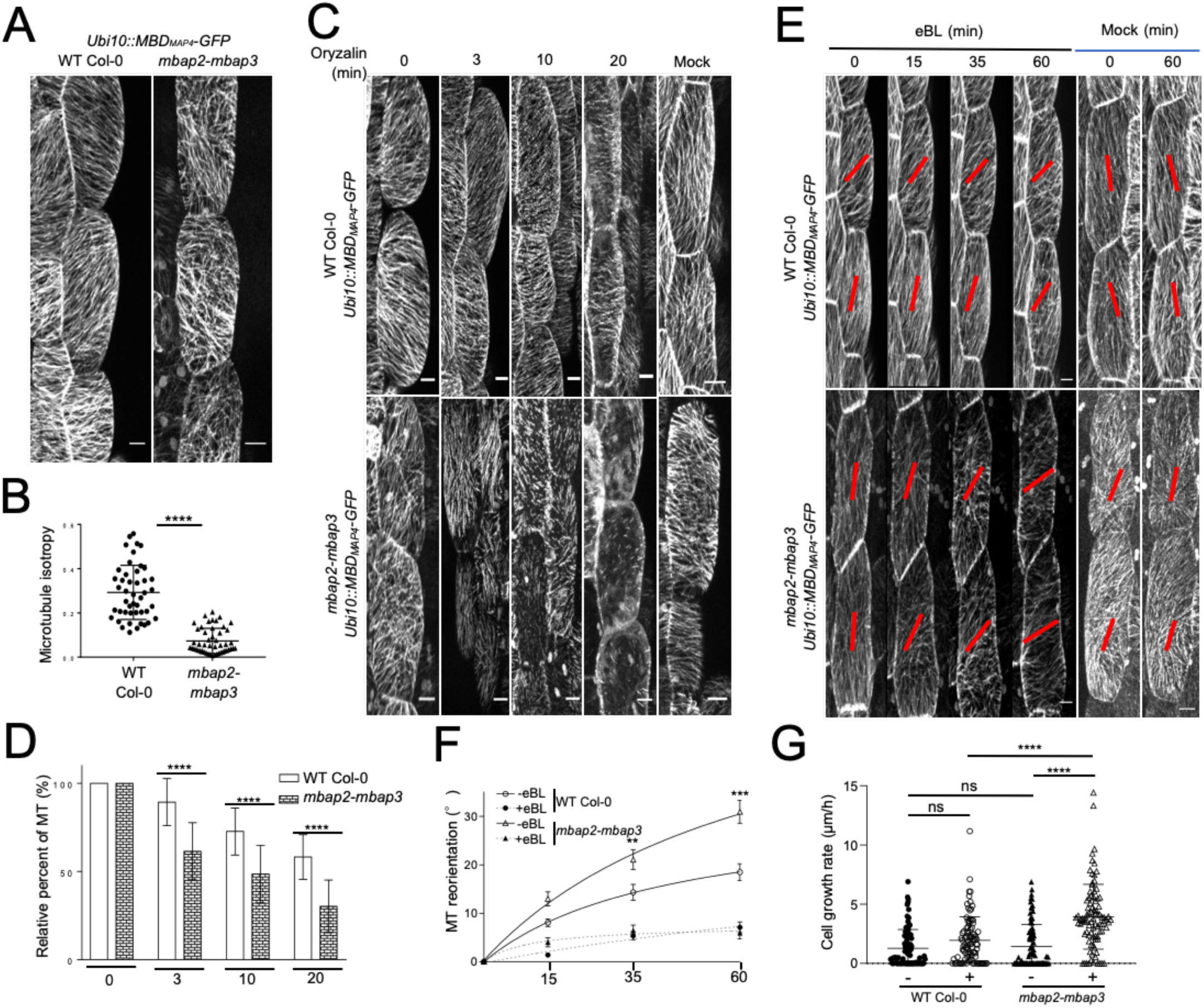
MBAP2/3 stabilize microtubules and suppress BR-induced microtubule reorientation in hypocotyls. (A) Cortical microtubule organization observed in 5-day-old Arabidopsis hypocotyl epidermal cells in WT and *mbap2-mbap3* plants expressing the *pUbi10::MBDMAP4-GFP* microtubule marker. Representative pictures are shown as maximum projection. (B) Quantification of the microtubules isotropy of cells shown in (A) using the Fibriltool ImageJ plugin. Over 15 cells from 3 hypocotyls were analyzed and experiments were independently repeated three times. Error bars indicate SD. Asterisks represent statistical significant difference (p-value < 0.0001, t-test). (C) Cortical microtubule organization in 5-day-old hypocotyl epidermal cells from WT and *mbap2-mbap3* seedlings expressing the *pUbi10::MBDMAP4-GFP* microtubule marker and treated with 50 µM oryzalin. Z-stacks were taken at 0, 3, 10 and 20 min after the treatment. Representative pictures are presented as maximum projections of Z-stacks. (D) Quantification of data as show in (C) using ImageJ to count filaments crossing a fixed line of 10 µm perpendicular to the mean orientation of CMTs within the cell. 15 cells from 3 independent hypocotyls in each condition were analyzed. Experiments were carried out in triplicates. Relative percentage of CMTs compared to time 0 is shown. t-test has been performed for each time point. Error bars represent SD and asterisks significative differences (p-value < 0.0001). (E) Cortical microtubule organization in 5-day-old hypocotyl epidermal cells from WT and *mbap2-mbap3* seedlings expressing the *pUbi10::MBDMAP4-GFP* microtubule marker and treated with 1µM eBL or mock. Images were taken at 0, 15, 35, and 60 min after the treatment. Representative pictures are presented as maximum projections of Z-stacks (F) Quantification of data as shown in (E) using the Fibriltool Image J plugin. 30 cells from 3 individual hypocotyls have been analyzed at each time point in three replicates. Error bars represent SEM. Asterisks represent statistical significant difference (p-value < 0.01 (**), 0.001 (***) ; t-test). (G) Cell growth rate during 1h eBL or mock treatment. Cell growth rate was measured between 0 and 60min after 1µM eBL treatment in WT and *mbap2-mbap3* hypocotyls. 30 cells from 3 individual hypocotyls were analyzed. Error bars correspond to SD, and asterisks represent significant differences (p-value < 0.01 (**), 0.001 (***), 0.0001 (****) ; t-test). Scale bar: 10 µm.

### MBAPs impact on the interaction of BRI1 with tubulin

Since the BR transcriptional reprograming is not affected in *mbap2-mbap3,* the BR responses driven by MBAPs likely involve their interaction with BRI1 and non-genomic responses through CMT reorganization. BRI1 dynamics was previously reported to require CMTs (Bücherl et al., 2017; Ruan et al., 2018; Xin et al., 2020). To better understand the connection between BRI1 and CMTs, we first carried out immunoprecipitation of the functional BRI1-mCit fusion, stably expressed in transgenic plants under the control of *BRI1* promoter, coupled to mass spectrometry analyses. Proteins were considered as interacting with BRI1 when specifically identified in the BRI1-mCitrine immunopurified fraction with at least two different peptides. Consistently, BRI1 was identified as the top hit in BRI1-mCitrine immunoprecipitates. Surprisingly, several peptides corresponding to α and Δ tubulin isoforms were found among the top hits (rank 5-9) (Supplemental Table 1). To confirm these observations, we carried out immunoprecipitation of BRI1-mCit from transgenic plants and immunoblotted against endogenous tubulin. We chose the MT marker GFP-MBD_MAP4_ as positive control for interaction with tubulin, and the generic PM reporter protein GFP-LTI6B (Shibasaki et al., 2009) as negative control. Although GFP-LTI6B strongly accumulated in inputs compared to BRI1, and thus was found at much higher levels after immunoprecipitation, tubulin was strongly enriched in the BRI1 complex (Fig. 6A, 6B). These findings support the scenario where BRs acts on CMTs directly through their association with BRI1. We next assessed whether MBAPs are bridging the interaction between BRI1 and CMTs. We therefore generated a *mbap2-mbap3*/BRI1-mCitrine transgenic line to evaluate the impact of the loss of MBAP2 and MBAP3 on the BRI1-tubulin interaction. Loss of MBAP2 and MBAP3 reproducibly yielded stronger immunoprecipitation of BRI1, even though the difference appeared non-significant due to the variability of the increase observed (Fig. 6C, 6D). These observations suggest that MBAPs likely inhibit the BRI1-tubulin association. Considering that BRI1 recycling from endosomes to the PM was previously reported in roots to depend on CMTs (Ruan et al., 2018), we tested if MBAPs were involved in BRI1 dynamics. No significant difference in BRI1 localization nor sensitivity to the endosomal-aggregating fungal drug BFA could however be observed (Supplemental Fig.S8).

**Fig 6.**
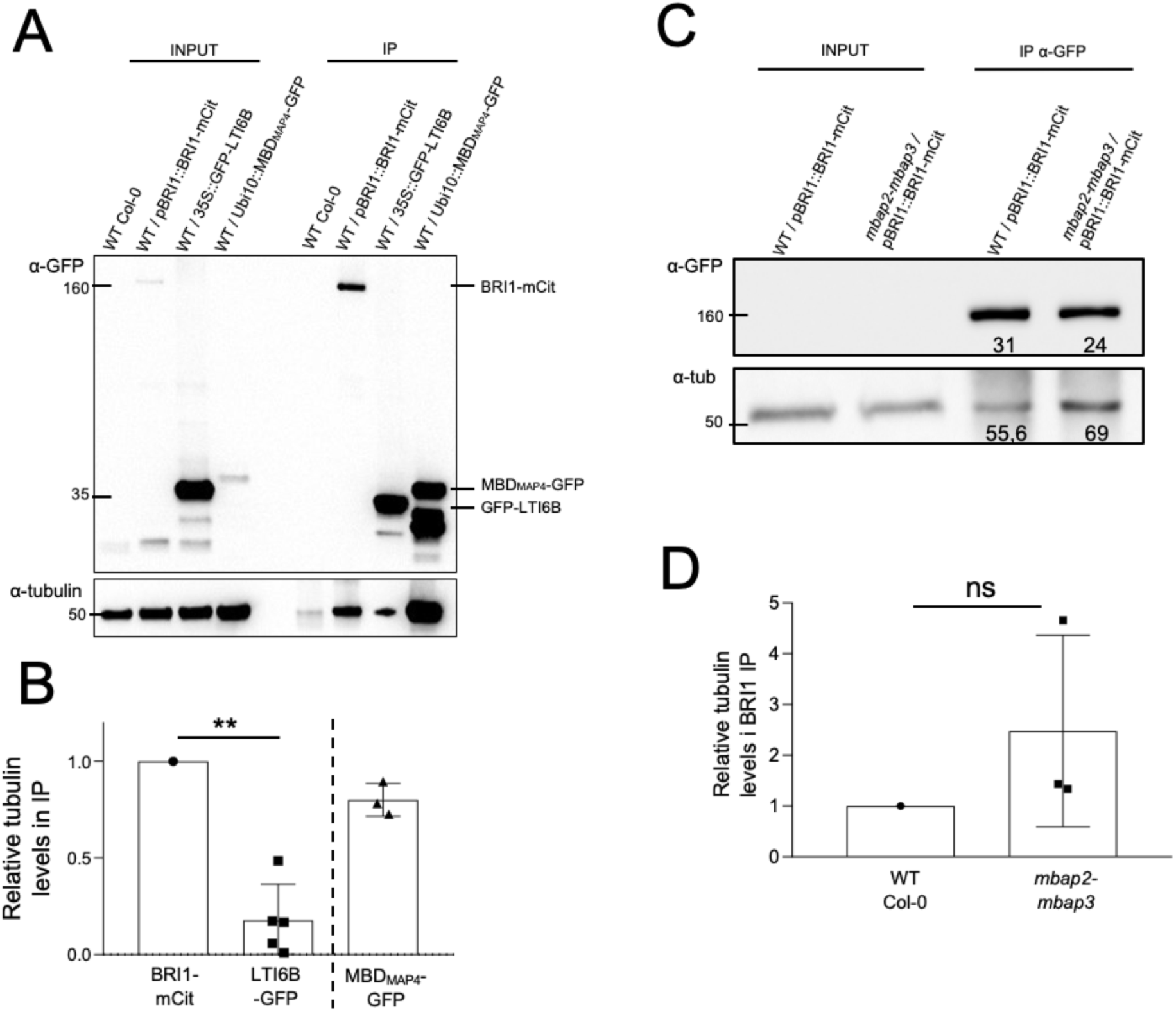
MBAP2/3 limit the interaction between BRI1 and tubulins. (A) Coimmunoprecipitation of α-tubulin with BRI1-mCit. 7-day-old WT, pBRI1::BRI1-mCitrine, p35S::GFP-LTI6B (negative control), or pUbi10::MBD_MAP4_-GFP plants (positive control) were subjected to IP using anti-GFP antibodies. Inputs (microsomal extract for WT, pBRI1::BRI1-mCitrine and p35S::GFP-LTI6B and soluble protein extract for pUbi10::MBD_MAP4_-GFP) and IP fractions were immunoblotted with antibodies against α-GFP and α-tubulin. (B) Ratio of coimmunoprecipitated α-tubulin relative to immunoprecipitated BRI1-mCit or control proteins. Experiments were repeated at least three times. Error bars indicate SD. Asterisks represent statistical significant difference (p-value < 0.01 (**) ; Mann-Whitney test) (C) Coimmunoprecipitation of α-tubulin with BRI1-mCitrine in 7-day-old WT and *mbap2-mbap3* expressing *pBRI1::BRI1-mCitrine.* A representative blot is shown. (D) Ratio of coimmunoprecipitated α-tubulin over immunoprecipitated BRI1-mCit. Experiments were done in triplicates.

## Discussion

CMTs have long been known to reorganize in response to various cues including phytohormones and environment signals, making them critical in plant development and adaptation (Chen et al., 2016; Hashimoto, 2015). However, the molecular mechanisms underlying CMT organization and reorientation remain unclear. Key players are MAPs that are known to regulate MT properties such as polymerization, bundling, severing or attachment to the PM (Hamada, 2014). Past work led to the identification of 727 MAPs in Arabidopsis, much of which have yet to be characterized (Hamada et al., 2013). Here, we uncovered a new family of MAPs involved in the transversal reorientation of CMTs induced by BRs in hypocotyls.

Several published evidences corroborate the fact that MBAPs act on CMTs and BR responses. The 3 MBAPs were found to be enriched in microtubule fractions, similar to their paralog CIP7 (Hamada et al., 2013), and thus belong to the Arabidopsis 727 MAPs. Recent work on CIP7 also confirmed its association to MTs and roles on CMT reorganization in response to light (Arico et al., 2024). Although the subcellular localization of the MBAP orthologs in rice (DEP2/EP2/SRS1/RELA) did not reveal MT association (Abe et al., 2010; Li et al., 2010; Zhu et al., 2021, 2010), a connection with BRs was proposed since DEP2/RELA interacts with the OsBZR1-interacting protein OsLic that activates BR transcriptional reprogramming during leave morphogenesis (Wang et al., 2008; Zhu et al., 2021). Additionally, the 3 MBAPs have been identified as partners of BIN2 through a proximity labelling approach. Phosphoproteomic analyses after treatment with the bikinin BIN2 inhibitor allowed the identification of the 3 MBAPs as phosphoproteins regulated by BIN2 (Kim et al., 2023). Our work now adds another layer to the link between MBAPs, CMTs and BRs with their identification as BRI1 partners and substrate, their CMT localization and their role as negative regulators of BR-mediated hypocotyl elongation.

MBAPs were identified through a yeast two hybrid screen using BRI1 kinase domain as bait, adding up to the long list of BRI1-associated proteins. Although we confirmed the MBAP1-BRI1 interaction by several independent approaches, we could not identify MBAPs in our mass spectrometry analyses of the BRI1 complex using plants grown in the absence of BR exogenous treatment. Since the MBAP-BRI1 interaction is not dependent on BRs, the absence of MBAPs in BRI1 immunoprecipitates rather suggests that MBAPs proteins are low abundant proteins. Regardless, MBAP proteins are not only interacting with BRI1 but also emerge as new BRI1 substrates, as evidenced by our *in vitro* kinase assays. This suggests that phosphorylation of MBAP1 occurs upon BRI1 activation following BR binding. This is however not accompanied by a dissociation of MBAP1 and BRI1 in our experimental conditions, in contrast to what has been described for other BRI1 partners BKI1 and BSKs (Wang and Chory, 2006; Tang et al., 2008).

MBAPs interact with BRI1 at the PM and in filaments that we characterized as CMTs, and where MBAP proteins are found. However, smaller and discontinuous filamentous structures were observed for the MBAP1-BRI1 interaction, in contrast to typical CMT pattern observed for reporter lines of MBAPs alone, which make us speculate on their nature. Since CMTs are known to be attached to the PM by several proteins such as CELLULOSE SYNTHASE-INTERACTIVE PROTEIN 1 or CLASP (Ambrose and Wasteneys, 2008; Lei et al., 2012; Li et al., 2012; Ruan et al., 2018), these structures could be the portion of CMTs that are tightly associated to PM and where BRI1 is found. BRI may therefore serve as an anchoring point of CMTs to the PM. Since CMTs have been shown to allow insertion of membrane proteins through transport vesicles (Ambrose et al., 2013), it is also possible that the MBAP-BRI1 interaction highlights contacts between CMTs and BRI1 endosomes, although no clear colocalization of MBAPs with BRI1-positive endosome was observed. Future studies using higher resolution microscopy will be needed to clarify where MBAPs and BRI1 meet in the cell and at CMTs.

The mechanisms by which MBAPs associate with CMTs is also unclear. No domain conferring MT binding are currently known. Association to such structures require specific conformation and charges (Hashimoto, 2015). Because MBAP1 is a large and insoluble protein, we were unable to test its association with MTs *in vitro* using classical centrifugation assay. We thus created truncations of MBAP1 and could identify a smaller fragment of MBAP1 located between amino acids 190 and 489 as sufficient to bind CMTs *in vivo* but not *in vitro*. Interestingly, CIP7 association to MTs has been shown to depend on its C-Ter region, which is not conserved with MBAPs (Arico et al., 2024). This supports the idea that conformation rather than protein sequence itself allows MT association. We narrowed down the the MT-binding domain of MBAP1 to an helix of 12 amino acids that is both necessary and sufficient for recruitment to MTs. The fact that the region harboring the MT-binding domain of MBAP1 fails to bind MT *in vitro* experiments may be explained by the requirement for other proteins to bridge CMTs and MBAPs, the requirement for post translational modifications, or oligomerization. Future work will be necessary to decipher the precise mechanism of MBAP interaction with CMTs.

Reverse genetic approaches using *mbap* knock-out mutants highlighted the involvement of MBAPs in BR signaling and growth. Although single mutants did not show any phenotype in resting conditions or in response to eBL/PPZ, double *mbap* mutant combinations showed hypersensitivity to BRs in light-grown hypocotyl elongation assays. Using *mbap2-mbap3* as a reference double mutant to dig further into the role of MBAPs, we uncovered that MBAPs do not participate to the canonical BR signaling pathway. Despite the hypersensitivity of *mbap2-mbap3* to BRs, no change in the typical and widely accepted BR response readouts were observed. Neither BES1 phosphorylation status nor expression of BR-regulated genes was affected in *mbap2-mbap3*. However, a clear disorganization of CMTs in hypocotyl cells and a strong hypersensitivity to oryzalin was observed in *mbap2-mbap3,* making these proteins stabilizer of the CMT network. Most importantly, loss of MBAP2 and MBAP3 yielded a faster and stronger reorientation of CMTs, pointing to MBAPs as negative regulator of the BR-mediated reorientation of CMTs. The enhanced elongation of hypocotyl cells observed in response to BRs in the *mbap* mutants supports the negative roles of MBAPs in BR-mediated hypocotyl growth. Reorientation of CMTs by BRs has been primarily shown to be induced through the transcriptional modelling of MAPs such as MDP40 and CLASP (Wang et al., 2012; Ruan et al., 2018). Here, we rather hypothesize that this is a direct effect through BRI1-MBAPs but the precise mechanism and contribution of the BRI1-MBAPs pairs remain to be clarified. BR perception by BRI1 likely inhibits MBAP activity through phosphorylation to allow CMT reorientation upon BR perception. This does not involve the dissociation of MBAPs from CMTs, but may necessitate a change in MBAP conformation or activity as shown for the CIP7 homolog (Arico et al., 2024), or partner proteins. The fact that BRI1 interacts with tubulins and that their association appears enhanced when MBAPs are missing suggests that CMT reorientation upon BR perception is in part driven by BRI1 interaction with tubulins and that must also involve still uncharacterized players. Therefore, inhibition of this association by MBAPs corroborates its repression of BR-mediated hypocotyl growth.

Based on all our findings, we established a hypothetical model of a parallel pathway contributing to the BR-dependent promotion of hypocotyl cell elongation through MBAPs (Fig. 7). In the absence of BRs, MBAPs interact with BRI1 at the PM and CMTs are stabilized. When cells are exposed to BRs, MBAPs are phosphorylated by BRI1 allowing the inhibition of MBAP function on CMTs, thus enhancing the BRI1/tubulin association. This leads to the reorientation of CMTs transversally to the growth axis although the precise events are still to be elucidated. CSC complex are thus running along the PM perpendicularly to the growth axis to release the brakes on cell elongation. Parallel to the MBAP-dependent pathway, the canonical BR signaling pathway ensures transcriptional reprogramming of the cell and allows remodeling of the CW and growth promotion. The ability of BIN2 to interact with MBAPs suggests that BIN2 may also contribute to the MBAP-mediated regulation of CMTs, also the mechanisms and contribution remains unknown.

**Fig 7.**
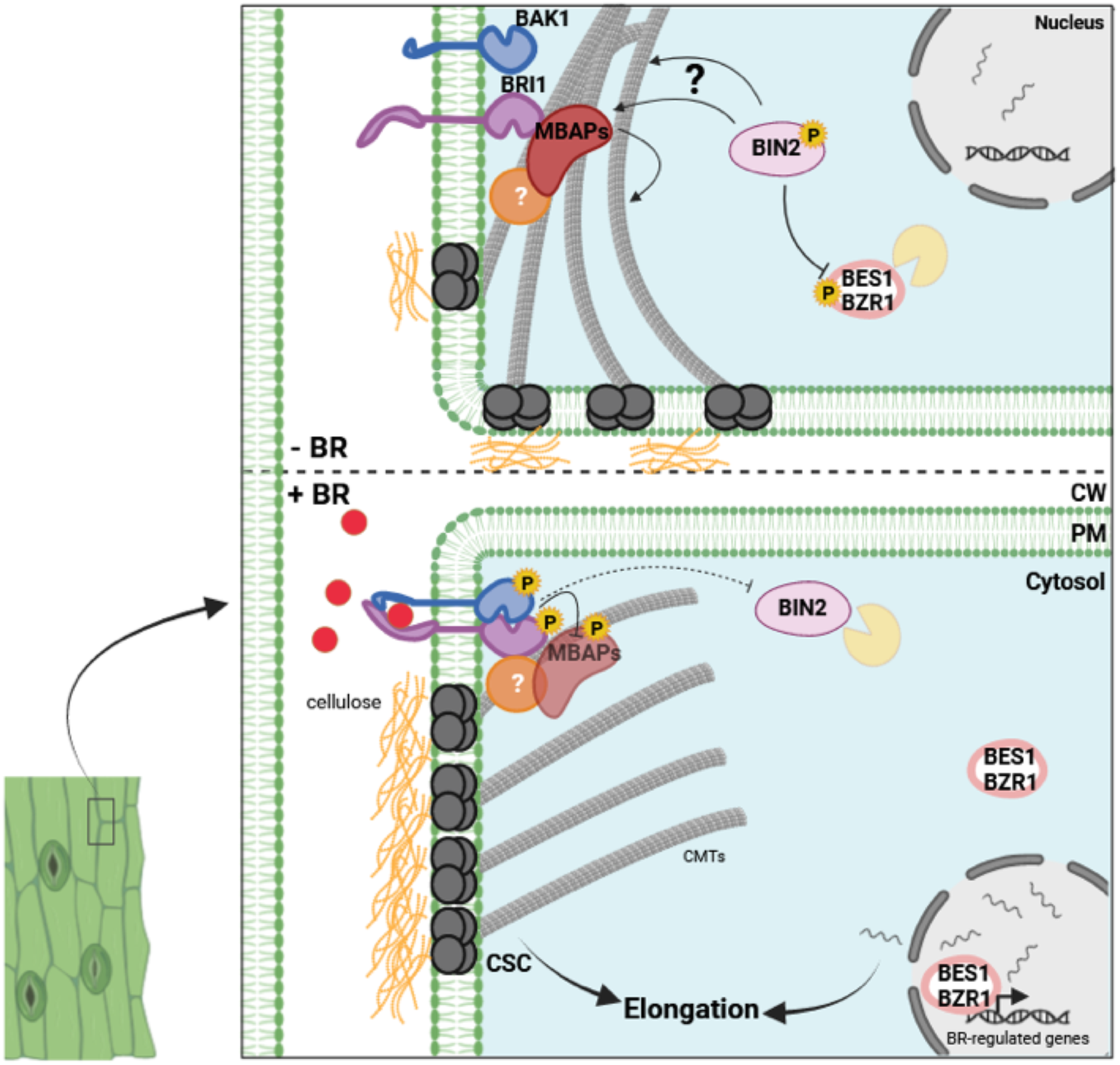
Hypothetical roles of MBAPs in BR-mediated hypocotyl elongation. In the absence of BRs, MBAPs stabilize CMTs which modulate insertion and mobility of the cellulose synthase complex (CSC). Cellulose is deposited equally in CW allowing growth to be isotropic. In response to BRs, the BRI1/BAK1 complex is activated, leading to BIN2 degradation and BES1/BZR1 dephosphorylation. The well-known transcriptomic changes associated with BR perception drive cell elongation. Parallel to this, we propose that BRI1 phosphorylates MBAPs which allows CMTs reorientation and strengthening CW perpendicularly to the growth axis. The reorientation mechanism must involve still unknown actors between MBAPs/CMTs and BRI1/CMTs. The canonical BR signaling pathway and the BRI1/MBAPs/CMTs pathway act together in the BR-mediated hypocotyl elongation. This figure was created using BioRender.

Altogether, our work unraveled a new non-genomic branch of the BR pathway that is required for proper cell elongation and plant growth. More work will be needed in the future to pinpoint the precise and respective biological roles of MBAPs, their regulation and the mechanisms governing their association to CMTs and BRI1. Besides, MBAPs may convey additional roles in the BR pathway considering their ability to interact with BRI1, BIN2 and maybe transcription factors (in rice) (Kim et al., 2023; Zhu et al., 2021). MBAPs could also serve as scaffolds as it has been shown for BSK3 and TETRATRICOPEPTIDE THIOREDOXIN-LIKE (Amorim-Silva et al., 2019; Ren et al., 2019). Especially the fact that TTL3 localizes to CMTs suggests possible functions of CMTs in this process (Xin et al., 2020). Finally, since BRI1 arose in monocotyledons (Kim and Russinova, 2020) and CMTs involve in a multitude of signal responses, MBAPs could act as bridge between membrane proteins and CMTs in other pathways providing faster adaptation to plants.

## Methods

### Plant materials and growth conditions

Arabidopsis Col-0 ecotype was used in all experiments. Arabidopsis T-DNA insertion lines for the three *MBAP* genes (Supplemental Table 2) were obtained from the Nottingham Arabidopsis Stock Centre (NASC). Genotyping was carried out as previously described (O’Malley et al., 2015), using primers as listed in Supplemental Table 2.

For stable Arabidopsis transgenic lines, WT*/*pBRI1::BRI1-mCitrine was already available in the lab (Martins et al., 2015). WT/pUBI10::GFP-MBD_MAP4_ were kindly provided by Dr. Geoffrey Wasteneys (University of British Columbia, Vancouver, Canada) (Ruan et al., 2018). WT/35S::GFP-LTI6B was provided by Dr. Ykä Helariutta (University of Helsinki, Finland) (Cutler et al., 2000). pBRI1::BRI1-mCitrine or pUbi10::GFP-MBD_MAP4_ markers were introgressed into *mbap2*-*mbap3* by crossing. *Ubi10::eGFP-MBAP1* or *pMBAP1-3::eGFP-GUS* constructs were transformed into WT or *mbap2 mbap3* by floral dip method with *Agrobacterium* GV3101 (MP90) strain (Clough and Bent, 1998). Monoinsertional homozygous transgenic lines were isolated.

Arabidopsis seeds were sterilized with a solution containing 12.5% bleach and 37.5% ethanol solution followed by brief washing with water, and stratified for 72h at 4°C on solid 0.8% agar (Kalys, HP 696) plates containing half-strength Linsmaier and Skoog (LS) (Caisson labs, LSP03) medium without sucrose. After stratification, plates were incubated in growth chambers under long-day conditions (16 h light/8 h dark, 90µE m^−2^s^−1^) at 21°C vertically for 5-7 days, depending on experiments.

For hypocotyl growth assays, Arabidopsis seedlings were grown on half-strength LS medium without sucrose supplemented with the corresponding concentrations of epibrassinolide (eBL, Sigma #E1641) or propiconazole (PPZ, Sigma-Aldrich # 45642). Hypocotyl length from 7-day-old seedlings was measured using the Fiji image software (http://fiji.sc/). For all growth parameters, at least 30 seedlings were used per each condition. Experiments were repeated three times.

### Gene cloning and vector construction

All cloning was done by multisite gateway technology (Thermo). All full length and partial coding sequences or promoter fragments were amplified by PCR from WT cDNA or genomic DNA using specific primers (Supplemental Table 3) and inserted into corresponding gateway entry vectors by BP reaction. Subsequently, fragments were combined with corresponding promoters, fluorescent proteins or tags into destination vectors by LR reaction (Marquès-Bueno et al., 2016) as listed in Supplemental Table 4. 35S::MBAP1_ΔVK_-mCherry was obtained by insertion of Bam*HI* restriction sites flanking the 36bp sequence encoding the VK helix by PCR (Supplemental Table 3), digestion using Bam*HI* and ligation. The *pBRI1::BRI1-mCit* construct was already available in the lab (Martins et al., 2015). The final constructs *35S::GFP-MBD_MAP4_* (Marc et al., 1998), *Ubi10::MBD_MAP4_-mCherry* (Ivanov and Harrison, 2014) were respectively gifted from Dr. Daniel Van Damme (VIB, Ghent) and Dr Denise Arico (LRSV, Toulouse). Vectors for production and purification of BRI1 cytoplasmic domain (amino acids 815-1196) pT7::8xHis-BRI1cp and its kinase-dead version pT7::8xHis-BRI1cp(D1027N) (Bojar et al., 2014) were given by Dr Michael Hothorn (University of Geneva, Switzerland). LecRK-I.9 in pDONR221 (Bouwmeester et al., 2011), BRL1/BRL3 in pDONR Zeo, and eGFP in pDONR221 or in pDONR-P2RP3 were respectively provided by Dr. Hervé Canut (LRSV, Toulouse), Dr Youssef Belkhadir (GMI, Austria) and Dr. Yoshihisa Oda (Nagoya University, Japan).

### Yeast two-hybrid screen

BRI1 cytoplasmic domain (amino acids 815-1196) was amplified using primers shown in Supplemental Table 3 and inserted into pDONR221 by BP reaction. The yeast two hybrid screen was performed by Hybrigenics (https://www.hybrigenics-services.com/).

### Chemicals

Chemical stock solutions were prepared in the following concentrations: 6 mM eBL (Sigma #E1641) in DMSO, 1 mM PPZ (Sigma) in water, 100 mM cycloheximide (Sigma # C7698) in ethanol, 10 mM BFA (Sigma # B7651) in DMSO, and 10 mM oryzalin (Sigma # 36182) in ethanol. The final concentrations of chemicals are indicated in the figure legends.

### GUS assay

GUS activity in seedling expressing *pMBAP1::eGFP-GUS, pMBAP2::eGFP-GUS, pMBAP3::eGFP-GUS and pBRI1::eGFP-GUS* was assayed using 5-bromo-4-chloro-3-indolyl β-D-glucuronide (X-Gluc) as substrate. 5-day-old seedlings were incubated in GUS staining solution (50 mM sodium phosphate pH 7.0, 0.1% Triton X-100, 0.5 mM K3/K4 FeCN, 1 mM X-Gluc) at room temperature for 8h. After rinsing in 70% ethanol for 1h, pictures were taken by light microscopy (Zeiss Axiozoom).

### Recombinant protein purification

*E Coli* BL21 strain DE3 carrying pT7::8xHis-BRI1cp or pT7::8xHis-BRI1cp (D1027N) were cultured in 1L of liquid LB media until OD600 reaches 0.6. Protein expression was induced by adding 0.5 mM IPTG for 16h at 16°C. Cells were collected by centrifugation (5,000 *xg*, 4°C, 1h) and resuspended in cold lysis buffer (20 mM Tris-base pH 8.0, 500 mM NaCl, 4 mM MgCl_2,_ 2 mM 2-Δ-mercaptoethanol, 1 mM Pefabloc (Merck 11429868001) and frozen in liquid nitrogen. After thawing, the pellet was lysed by adding 0.5 mg/mL lysozyme and incubated for 45 min on ice before 8 cycles of 45 seconds (50 kHz) sonication. The lysate was centrifuged (7000 x*g*, 4°C, 1h), and the supernatant was recovered. His-tagged proteins were purified using HisTrap HP Ni^2+^ affinity column (GE Healthcare 17-5255-01) according to the manufacturer’s instruction. Briefly, the supernatant was applied onto equilibrated resin in a column, and the resin was washed four times with 20 mL lysis buffer by increasing the concentration of imidazole from 10, 30, 50, 75 mM in each wash. Then, His-tagged proteins were eluted using lysis buffer supplemented with 200 mM imidazole. The collected samples were mixed with glycerol at 10% final concentration and stored at -80°C. For MBP-tagged protein expression, BL21 RiPL codon plus harboring corresponding plasmids were grown in 1L liquid LB supplemented with glucose (2 g/L). Protein expression was induced with 0.3 mM IPTG for 1h at 37°C (MBP alone) and for 6h at 19°C (MBP-MBAP1 full length and fragments) after OD600 reached 0.7. Cells were collected by centrifugation (4000 *xg*, 4°C, 30min), and the pellet was resuspended in cold lysis buffer (20 mM Tris-base pH 7.4, 200mM NaCl, 1 mM EDTA, 1 mM DTT, 1 mM Pefabloc, 1 mM PMSF) followed by 8 cycles of 15s (50 kHz) sonication. Supernatant was recovered after centrifugation (12000 x*g*, 4°C, 1h) and incubated with 0.002% equilibrated amylose resin (NEB) for 2h on a rotating wheel at 4°C. Beads were collected by centrifugation (500 *xg*, 4°C, 10 min) and washed four times in fresh lysis buffer for 10 min at 4°C. Proteins were eluted using the lysis buffer containing 10 mM maltose and mixed with glycerol to a final concentration of 10% for storage at -80°C. Purified proteins were checked by western blotting with anti-His-tag or anti-MBP antibodies. The fractions displaying the cleanest (visualized by Coomassie staining) and the highest levels of purified proteins (measured by Bradford assay) were used for in vitro assays.

### In vitro kinase assay

500 ng of kinase (His-cpBRI1 or His-cpBRI1 kinase-dead) were incubated with 2 µg of substrates (MyBP (Myelin basic protein), MBP, or MBP-MBAP1) in a 20µL kinase reaction buffer (20 mM Tris-HCl pH 7.5, 10 mM MgCl_2_, 100 mM NaCl, 1 mM DTT, 100 µM ATP) for 3 hours at 30°C. Reactions were stopped by adding 2x Laemmli buffer (125 mM Tris base, 4% SDS, 20% glycerol, 10% 2-ϕ3-mercaptoethanol, 0.001% bromophenol blue) and boiling for 10 min at 95°C. Samples were analyzed by western blotting using anti-phospho-threonine antibodies.

### MT sedimentation assay

In vitro microtubule binding assay was carried out with Microtubule Binding Protein Spin-Down Assay Biochem kit (Cytoskeleton BK029) by following the manufacturer instructions. Bovin Serum Albumin (BSA) and MAP fraction were used as negative and positive controls, respectively. After separating proteins on acrylamide gels, proteins were stained by Coomassie staining.

### Nicotiana benthamiana infiltration

For transient expression in *N. benthamiana* leaves, *Agrobacterium tumefaciens* (GV3101 MP90) was transformed with corresponding plasmids and cultured in liquid medium with antibiotics. After collecting pellets, bacterial pellets were adjusted to OD600 of 0.1 (MAP4 constructs) or 0.5 (all other constructs) with infiltration buffer (10 mM MES pH 5.6, 10 mM MgCl_2_, 100 µM Acetosyringone). For coexpression analysis, Agrobacterium solutions harboring different plasmids were mixed in a 1:1 ratio [v:v]. All combinations were coinfiltrated with the silencing suppressor p19 (Voinnet et al., 1999) into 4-5-week-old plants. Leaf discs were analyzed 2 days after infiltration under the confocal microscope. For coimmunoprecipitation, 2 day-postinfiltration leaves were treated by infiltration with 10 µM eBL or mock solution for 20 min before freezing in liquid nitrogen.

### Coimmunoprecipitation

Coimmunoprecipitation from microsomal fractions was performed using 1 g of infiltrated *N. benthamiana* leaves or 1 g of 7-day-old Arabidopsis seedlings, depending on the experiment. Microsomal protein fractions were obtained as previously described (Jaillais et al., 2011). Immunoprecipitation was carried out using micromagnetic bead-coupled GFP antibodies from µMACS GFP Isolation Kit (Miltenyi Biotec 130-091-125) following the manufacturer’s instruction. Immunoprecipitated and microsomal fractions were analyzed by western blotting. For GFP-MBD_MAP4_, IP were performed directly on total proteins without microsomal extraction. Experiments were done at least in triplicates.

### IP MS/MS

Immunoprecipitation-coupled MS/MS analyses were performed using 1 g of 12-day-old light-grown *pBRI1::BRI1-mCit* or wild-type seedlings. Microsomal fractions were obtained as described above. Immunopurified proteins were reduced with dithiothreitol, alkylated with iodoacetamide and digested overnight with trypsin. Salts and reagents were removed using reversed phase (C-18) cartridge clean-up. The salt-free peptides were dissolved in 0.1% formic acid and subjected to ESI-MS/MS analysis on a Thermo LTQ-Orbitrap XL instrument. A capillary column (inner diameter 75 µm, packing length 10 cm of C-18 silica) with an integrated spray tip was used at a 300 nL/min 0.1% formic acid/acetonitrile gradient. Peptide precursor masses were determined with high accuracy by Fourier transform MS in the Orbitrap followed by data-dependent MS/MS of the top five precursor ions in each chromatographic time window. Data were analyzed using the Mascot algorithm (Matrix Science, London, UK on a local Mascot server (version 2.1.0) and searched against the latest Swiss Protein database. Hits were defined as i) found in coimmunoprecipitates with BRI1-mCit and absent from wild-type, and ii) identified with at least 2 unique peptides.

### Protein extraction

For detection of BES1, 100 mg of 7-day-old Arabidopsis seedlings were treated with mock or 1µM eBL for 20 min and frozen in liquid nitrogen. Total proteins were extracted using Laemmli 4X extraction buffer (250 mM Tris base, 8% SDS, 40% glycerol, 20% 2-mercapto-ethanol, 0.002% bromophenol blue) at a 1:3 [w/v] ratio between tissue powder and buffer. After centrifugation (10,000 *xg*, 5 min), the supernatant was recovered as a crude protein fraction.

### Western blot

Protein samples were denatured for 5 min at 95°C and separated in TGX Stain-Free FastCast 10% Acrylamide gel (Biorad #1610183) using running buffer (25 mM Tris base pH 8.3, 190 mM glycine, 0.1% SDS) at 130 V constant setting. Proteins were then transferred onto a nitrocellulose membrane 0.45 μm (Biorad Cat # 1620115) using a wet transfer system with modified Towbin transfer buffer (25 mM Tris base pH 8.3, 190 mM glycine, 20% ethanol). After blocking the membranes with 5% skim milk in TBS-T (50 mM Tris pH 7.6, 150 mM NaCl, 0.1% Tween-20) for 1h, membranes were probed overnight at 4°C with the corresponding antibodies : anti-GFP-conjugated with HRP (Miltenyi Biotec 130-091-833, 1:5000), anti-RFP (Abcam 34767, 1:5000), anti-αTubulin (Agrisera, AS10 680, 1:5000), anti-Phospho-threonine (Cell Signaling 9381S, 1:1000), anti-MBP (NEB E8038S, 1:2000) or anti-BES1(Yin et al., 2002) antibodies. All antibodies were diluted in 5% skim milk TBS-T. After washing the membranes three times with TBS-T for 15 min, membranes were probed for 1h at room temperature with the corresponding secondary antibody (anti-rabbit-conjugated with HRP (Agrisera AS09 602, 1:10000) or anti-rat-conjugated with HRP (Invitrogen 31470, 1:5000)) when necessary. Membranes were washed three times for 15 min with TBS-T followed by washing TBS for 5 min to remove Tween-20. Peroxidase activity was detected using the SuperSignal West Femto substrate kit (Thermo Scientific 34094) and a ChemiDoc Imaging system (Biorad). Coomassie-stained gels (Sigma 27813) or ponceau-stained membranes (Ponceau S 0.1%, acetic acid 5%) were used as loading control.

### Confocal microscopy and image quantification

For *N. benthamiana* leaves, leaf discs were mounted between slides and coverslips before imaging. Arabidopsis seedlings were treated as described in the legends to figures prior to mounting. For eBL treatment, seedlings were directly mounted in eBL or mock solution on slides. To visualize Arabidopsis cortical microtubule organization, images were taken on the proximal (upper part) of light-grown hypocotyls from 5-day-old seedlings. Images were captured under an upright Leica TCS SP8 confocal laser scanning microscope (www.leica-microsystems.com/home/). The excitation and detection wavelength were set to obtain signal as below (excitation, detection window); GFP (488 nm, 500-550 nm), mCit (514 nm, 530-575 nm), mChe (561 nm, 580-630 nm), and GFP/mCherry (488/561 nm, 500-550 nm, 580-630 nm with sequential scan). Laser intensity and detection settings were kept constant when required. Before imaging, care was taken to be below saturation levels. Z-stacks were taken with images every 1 µm and shown as maximum projection. Isotropy degree and mean orientation of MTs within a region of interest (one cell) were measured by the Fibriltool Image J plugin (Boudaoud et al., 2014). To quantify number of CMTs in a cell, the number of MTs crossing a fixed 10 µm line transversally to the mean orientation of CMTs were counted (Wang et al., 2012). To quantify BRI1 localization, mCitrine intensity of the whole cell and intracellular regions were scored using ImageJ, as already described for membrane proteins (Spielmann et al., 2023). At least 10 cells in 3 independent hypocotyls or roots were imaged in triplicates.

### Multiple sequence alignment

Searching gene orthologs of the MBAPs in the *viridiplantae* were carried out using PLAZA interface (https://bioinformatics.psb.ugent.be/plaza/versions/plaza_v5_dicots/ genes/view). Arabidopsis MBAP (AT1G17360.1, AT1G72410.2, AT3G14172.1) protein sequences were aligned using MUSCLE (https://www.ebi.ac.uk/Tools/msa/muscle/) (Edgar, 2004). Detection of conserved motifs in protein sequences between MBAPs, CIP7 and orthologs were carried out using the MEME tool (https://meme-suite.org/meme/tools/meme).

### RT-PCR

Total RNA was isolated using the RNeasy Plant Mini Kit (QIAGEN). RNA integrity was checked by agarose gel electrophoresis. Reverse transcription, digestion of genomic DNA, and cDNA synthesis were carried out using the RevertAid RT Reverse Transcription Kit (Thermo Scientific). PCRs were run with GoTaq DNA Polymerase (Promega) by following the instructions of the manufacturer. 25 or 30 cycles were set to amplify *ACT2,* and *MBAP*s and *BRI1* respectively with primers listed in Supplemental Table 5.

### Quantitative real-time PCR

RNA was isolated from BR-treated or non-treated seedlings and cDNAs prepared as described above. qPCR was done in a 15 µL final volume containing 10 ng cDNA, 0.3 µM of each gene-specific primer (Supplemental Table 6), and Maxima SYBR green qPCR (Thermo Fisher Scientific). Primer efficiency has already been validated (Barberon et al., 2011; Oh et al., 2012; Saeed et al., 2023; Spielmann et al., 2020). PCRs were performed using real-time PCR system CFX Opus 384.

### Statistical analysis

Statistical tests, number of biological replicates, samples in each biological replicate, and significance are mentioned in legends. All tests and graphs were performed using GraphPad Prism 7 software.

## Acknowledgements

We would like to thank scientists who kindly shared published materials. We also acknowledge the Imaging facility from the Fédération de Recherche Agrobiosciences Interactions et Biodiversité of Toulouse (FRAIB) for their help for imaging and image analyses. This work was supported by research grants from the French National Research Agency (ANR-22-CE13-0021-01 to S.F., ANR-17-CE20-0026-01 to G.V.) and the French Laboratory of Excellence (project “TULIP” grant nos. ANR–10–LABX–41 and ANR–11–IDEX–0002–02 to G.V.).

