## Supplemental Figures and Tables for "Direct non-transcriptional link between brassinosteroid perception and cortical microtubule reorientation drives hypocotyl growth"

### Figure S1

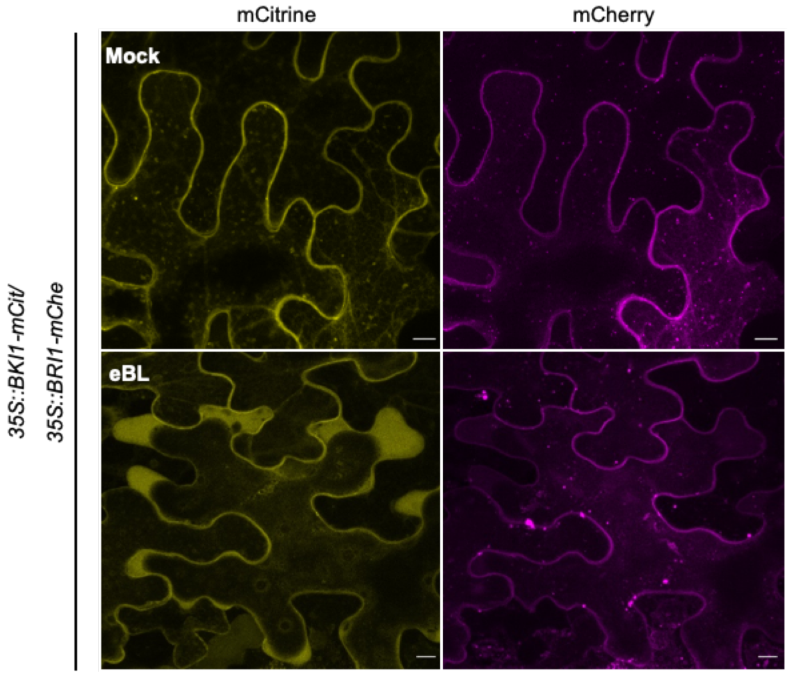


**Fig. S1. eBL treatment in *N. benthamiana* is sufficient to release BKI1 from the PM to the cytosol.** *N. benthamiana* leaf discs coexpressing *35S::BKI1-mCitrine* and *35S::BRI1-mCherry* infiltrated with or without eBL (10 µM) were observed after 20 min by confocal microscopy. Panels correspond to maximum projection of Z-stacks. Scale bar: 10 µm.

### Figure S2

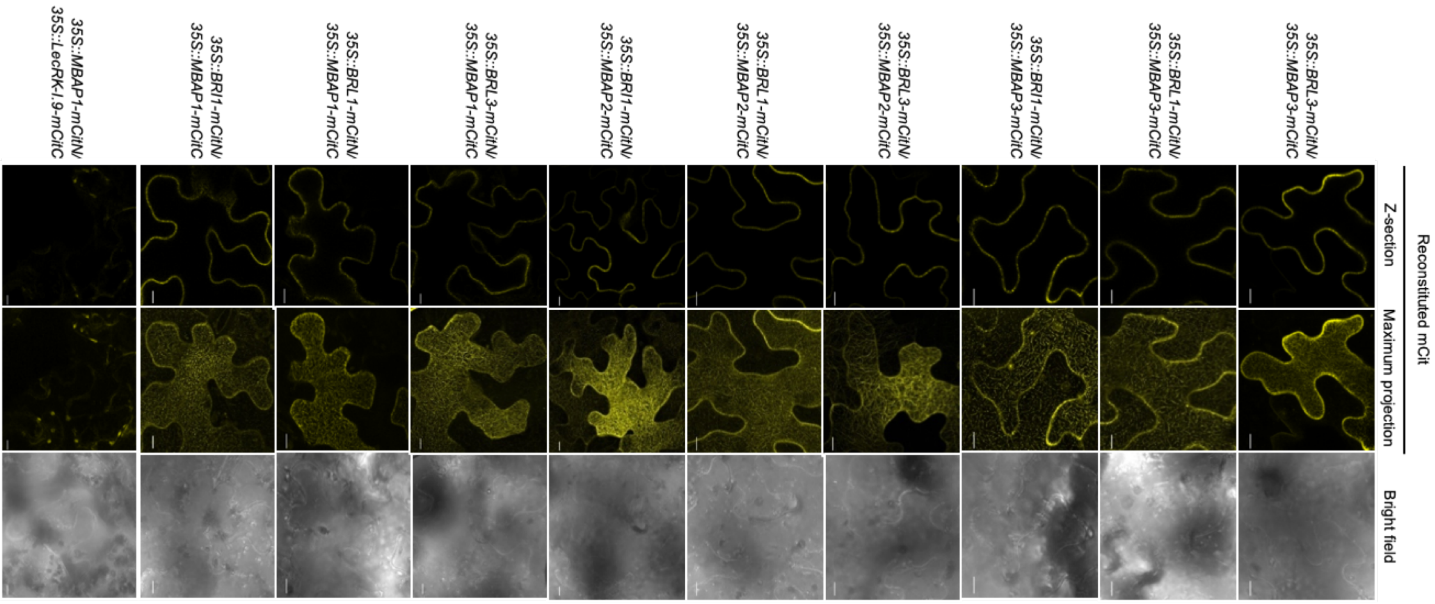


**Fig. S2. Interaction network between MBAPs BRI1/BRLs.** Bimolecular Fluorescence Complementation in *N. benthamiana* coexpressing *35S::BRI1-mCitN or 35S::BRL1-mCitN or 35S::BRL3-mCitN* and *35S::MBAP1-mCitC* or *35S::MBAP2-mCitC* or *35S::MBAP3-mCitC*. Each combination is indicated in the figure. The combination *35S::MBAP1-mCitC* with *35S::LecRK-I.9*-mCitC was used as a negative control.

### Figure S3

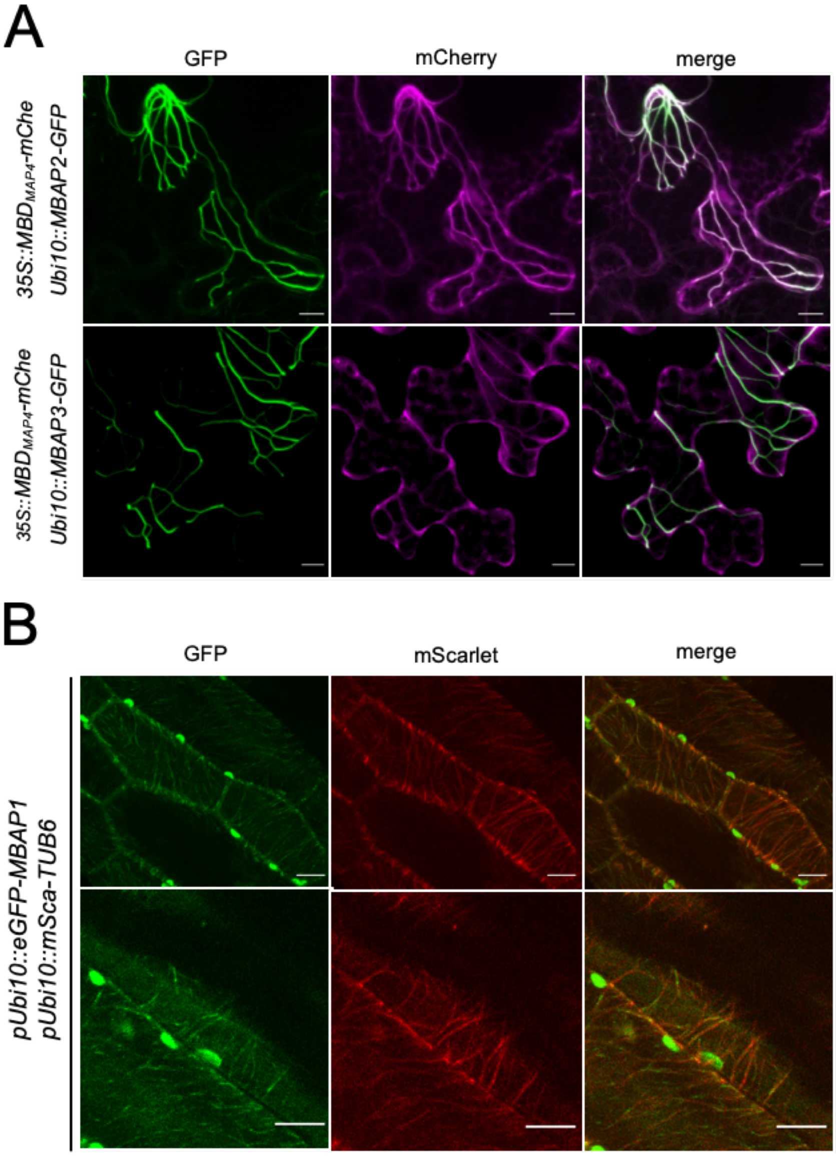


**Fig. S3. MBAP2 and MBAP3 localize to CMTs.** Representative pictures of *N. benthamiana* leaf epidermal cells coexpressing *pUbi10::MBAP2-eGFP* or *pUbi10::MBAP3-eGFP* and *pUbi10::MBDMAP4-mCherry*. Maximum projection of Z-stacks is shown. Scale bar: 10 µm. (B) Covisualization of pUBQ10::mEGFP-MBAP1 and pUBQ10::mScaI-TUB6 in hypocotyls from stable Arabidopsis transgenic lines.

Figure S4

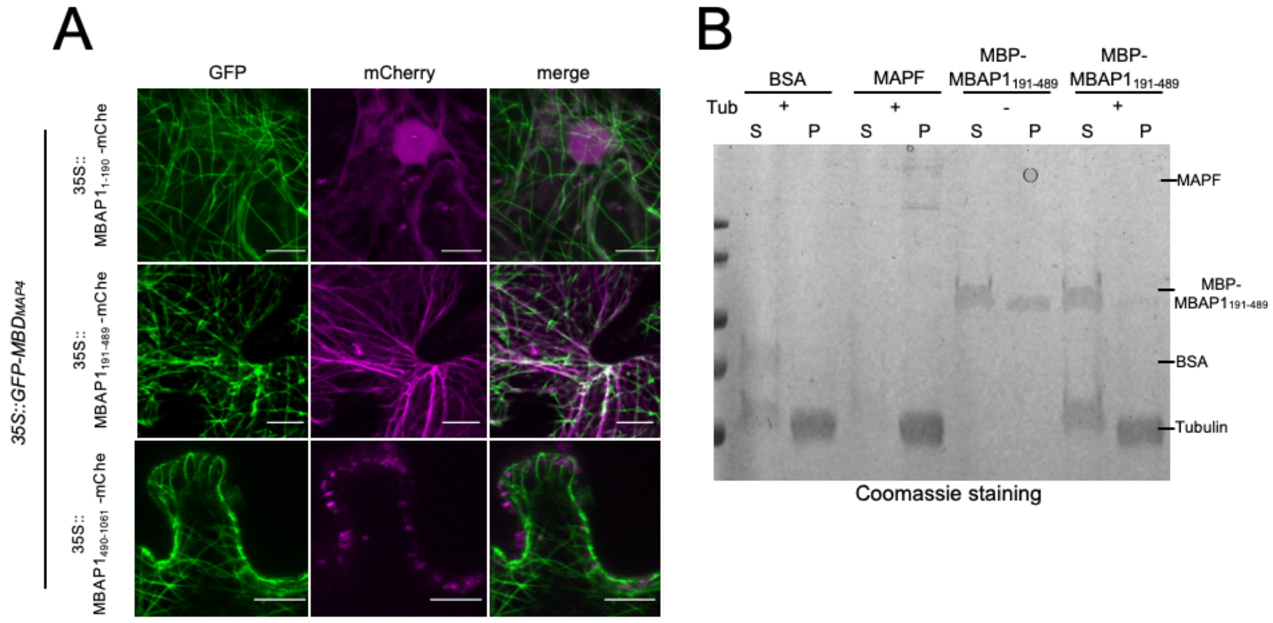


**Fig S4. The central region of MBAP1 is responsible for CMT association *in planta*. (A)** *N. benthamiana* leaf discs coexpressing *p35S::GFP-MBDMAP4* and the 3 fragments (aa 1-190, 191-489, 490-1061) from MBAP1 and fused to mCherry (*35S::MBAP1$_{1\text{-}190}$-mCherry*, *35S::MBAP1$_{191\text{-}489}$-mCherry*, *35S::MBAP1$_{490\text{-}1061}$-mCherry*) were observed by confocal microscopy. Panels are shown as maximum projection of Z-stacks. Scale bar: 10 µm. (B) *In vitro* microtubule centrifugation assay of 191-489MBAP1 fragment fused to MBP (MBP-191-489MBAP1) responsible for MT association *in planta*. BSA and MAPF (Microtubule associated protein fraction) were used as negative and positive controls, respectively.

### Figure S5

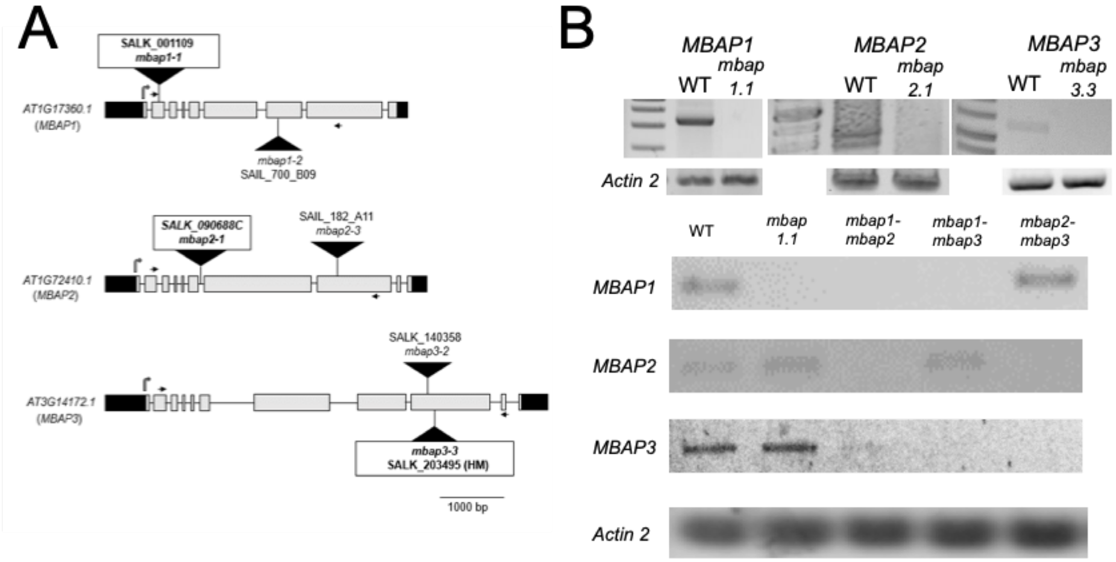


**Fig S5. Characterization of knockout mutants for *MBAP1*, *MBAP2,* and *MBAP3*.** (A) Insertions are indicated by a black triangle. The boxed insertion lines are used to generate the double mutants by crossing. The start codon is indicated by an upright grey arrow. 5' and 3' UTRs are represented as black boxes, and exons shown in light grey. Primers to genotype and to inspect gene expression are depicted as black arrows. Scale bar: 1000bp. (B) Expression levels of *MBAP1*, *MBAP2*, *MBAP3* in the corresponding single and double mutant combinations.

### Figure S6

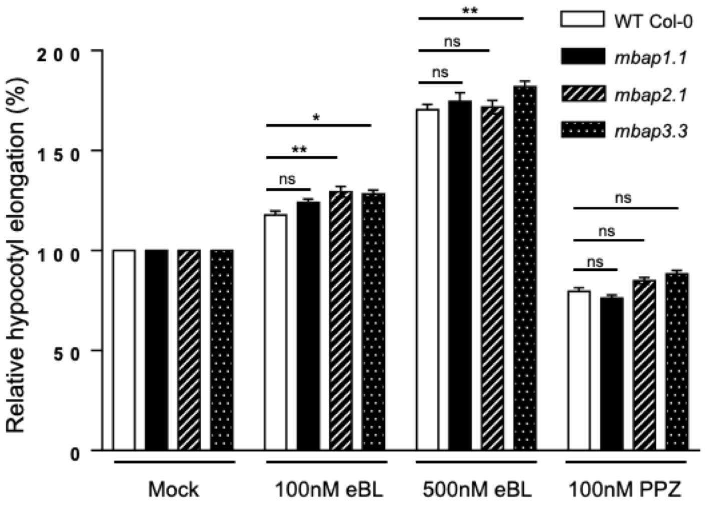


**Fig S6. Sensitivity of single knock out mutants to BR and PPZ.** Relative hypocotyl length of 7-day-old WT, *mbap1.1*, *mbap2.1,* and *mbap3.3* seedlings grown on half-LS media and containing eBL (100nM, 500nM) or propiconazole (100nM PPZ). Data are shown relative to mock. Experiments were carried out in triplicates. Error bars indicate SEM (n>22). Asterisks mean significant differences (p-value < 0.05 (*), 0.01 (**), and n.s. to no significant difference ; Two-way ANOVA with Dunnett's multiple comparison test).

### Figure S7

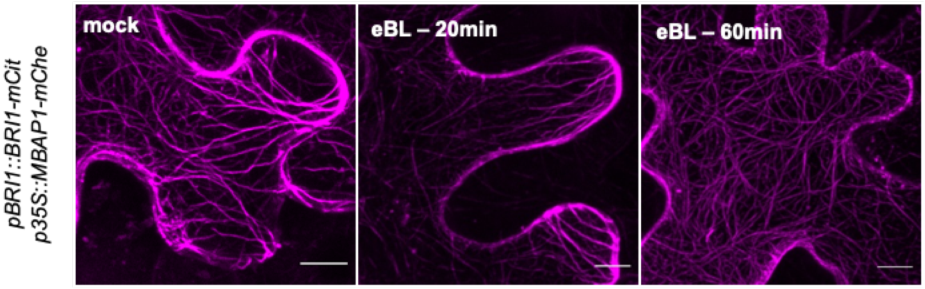


**Fig S7. BRs do not affect MBAP1 microtubule pattern in *N. benthamiana*.** *N. benthamiana* leaf discs coexpressing *pBRI1::BRI1-mCitrine* and *p35S::MBAP1-mCherry* were observed by confocal microscopy in mock condition or after infiltration of eBL (10 µM). Representative images are shown as maximum projection of Z-stacks. Scale bar: 10 µm.

### Figure S8

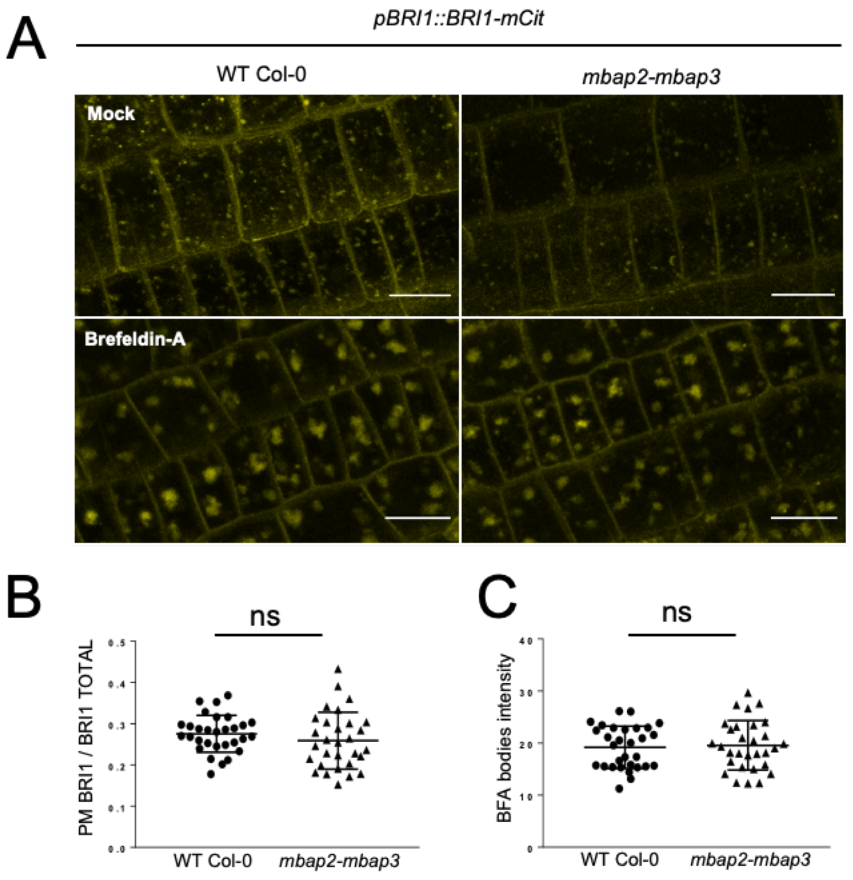


**Fig S8. Localization and recycling of BRI1 is not affected in *mbap2-mbap3* knock out mutant.** (A) Confocal microscopy observations of root epidermal cells of 5-day-old WT and *mbap2-mbap3* seedlings expressing *pBRI1::BRI1-mCitrine* in mock condition or cotreatment of 50 µM BFA/100 µM CHX for 30 min. Maximum projection of Z-stacks in the entire cells with 1 µm steps. Scale bar: 10 µm (B) Quantification of ratio between PM-localized BRI1 and total BRI1 signal intensity in WT and *mbap2-mbap3*. PM intensity was calculated by subtracting intracellular signal intensity from total signal intensity using Image J. (C) Quantification of BFA sensitivity by measuring mean intracellular fluorescence intensity. Symbols indicate each data point, and error bars are SD. Measurement was done in 10 cells from 3 independent roots showing no significant difference (t-test). Experiments were done in triplicates.

**Table S1. BRI1 interactors identified by IP/MS**

| Rank | Accession | Protein name | Maximum number of unique peptides |
|---|---|---|---|
| 1 | AT4G39400 | Brassinosteroid insensitive 1 (BRI1) | 12 |
| … | | | |
| 5 | AT1G50010 | Tubulin alpha-2/alpha-4 chain (TUA2) | 5 |
| 6 | AT5G62690 | Tubulin beta-2/beta-3 chain (TUB2) | 4 |
| 7 | AT5G12250 | Tubulin beta-6 chain(TUB6) | 4 |
| 8 | AT5G44340 | Tubulin beta-4 chain(TUB4) | 3 |
| 9 | AT1G20010 | Tubulin beta-4 chain(TUB5) | 3 |
| … | | | |

Among the 102 putative BRI1 interacting-proteins identified by coIP combined with mass spectrometry, only relevant proteins to the study are shown according to their TAIR annotation (The Arabidopsis Information resource ; https://www.arabidopsis.org).

**Table S2. T-DNA lines, insertion position, and primers used for genotyping**

| Line | Allele | Type | Insertions | Position | Primer T-DNA | Primer LP | Primer RP |
|---|---|---|---|---|---|---|---|
| SALK_001109 | *mbap1-1* | SALK | AT1G17360.1 | 2nd Exon | GCGTGGACCGCTTGCTGCAACT | TGCATGCTTCACTGATGAAAG | ATGGAGCTGCATTTTCATCTG |
| SAIL_700_B09 | *mbap1-2* | SAIL | AT1G17360.1 | 7th Exon | TAGCATCTGAATTTCATAACCAATCTCGATACAC | TACCATTTCTCGCCTGTGTTC | AAAGGCAGGAAGAAAGAGCTG |
| SALK_090688C | *mbap2-1* | SALK | AT1G72410.1 | 6th Intron | GCGTGGACCGCTTGCTGCAACT | GACGTTGCTTCAACTCCTGAC | ATTGTGATTCTTGACCGATCG |
| SAIL_182_A11 | *mbap2-3* | SAIL | AT1G72410.1 | 8th Exon | TAGCATCTGAATTTCATAACCAATCTCGATACAC | CTCCAGACGTGGTAAGTCGAG | TCCTCAAACACCAATGTTTCC |
| SALK_040438 | *mbap3-1* | SALK | AT3G14172.1 | 7th Exon | GCGTGGACCGCTTGCTGCAACT | TGCTGGTTTCAACCCTATCAC | CCACAGCTGTTTTCTCTCCAG |
| SALK_203495 | *mbap3-3* | SALK | AT3G14172.1 | 9th Exon | GCGTGGACCGCTTGCTGCAACT | TTTAAAGCTCAATGGTGTGCC | AAGAAGAAAAGTTGCGAAGGC |

**Table S3. Primers used for cloning DNA sequences in the corresponding pDONR entry vectors**

| DNA amplified | Primer F | Primer R | Vector |
|---|---|---|---|
| BRI1cp for Y2H screen | GGGGACAAGTTTGTACAAAAAAGCAGGCT GGAGAGAGATGAGGAAGAGACGG | GGGGACCACTTTGTACAAGAAAGCTG GGTATCATAATTTTCCTTCAGGAACTTC | pDONR221 |
| CaMV35S | GGGGACAACTTTGTATAGAAAA GTTGCTCGCGGCCAACATGGTGGA | GGGGACAACTTTGTATAGAAAAGTT GCTCGCGGCCAACATGGTGGA | pDONR-P4P1R |
| pUbi10 | GGGGACAACTTTGTATAGAAAA GTTGCTAGTCTAGCTCAACAGAGC | GGGGACTGCTTTTTTGTACAAACTT GCCTGTTAATCAGAAAAACT | pDONR-P4P1R |
| mCherry | GGGGACAACTTTGTATAATAAAGTTGCTT ACTTGTACAGCTCGTCCATGCCGCCGGTGGA | GGGGACAGCTTTCTTGTACAAAGTG GCTGTGAGCAAGGGCGAGGAG | pDPONR-P2RP3 |
| mCitrineN | GGGGACAGCTTTCTTGTACAAA GTGGCCATGGTGAGCAAGGGCGAG | GGGGACAACTTTGTATA ATAAAGTT GATTAGGCCATGATATAGACGTTGTGG | pDPONR-P2RP3 |
| mCitrineC | GGGGACAGCTTTCT TGTACAAA GTGGCCGACAAGCAGAAGAACGGCATC | GGGGACAACTTTGTATAATAAAGTT GATTACTTGTACAGCTCGTCCATG | pDPONR-P2RP3 |
| mCitrine | GGGGACAGCTTTCTTGTACAAA GTGGCTATGGTGAGCAAGGGCGAG | GGGGACAACTTTGTATAATAAAGTT GCTTACTTGTACAGCTCGTCCATGCCG | pDPONR-P2RP3 |
| MBAP1 CDS | GGGGACAAGTTTGTACAAAAAAGCAGGC TGGATGAAGGCTGATACTGTTCTAGAC | GGGGACCACTTTGTACAAGAAAGCT GGGTATCTTGGCTTCGAGTCGTTTCC | pDONR221 |
| MBAP1 stop CDS | GGGGACAGCTTTCTTGTACAAAGTGGCC ATGAAGGCTGATACTGTTCTA | GGGGACAACTTTGTATAATAAAGTTG ATTATCTTGGCTTCGAGTC | pDONR-P2RP3 |
| MBAP1 stop CDS | GGGGACAAGTTTGTACAAAAAA GCAGGCTGGATGAAGGCTGATACTGTTCTAGAC | GGGGACCACTTTGTACAAGAAAGCT GGGTAATTTCTTGGCTTCGAGTCGTT | pDONR221 |
| MBAP1 VK CDS | GGGGACAAGTTTGTACAAAAAA GCAGGCTGGATGGTACAAGATAGGATTA | GGGGACCACTTTGTACAAGAAAGCT GGGTACTTTTGTTTGCTCTCAAA | pDONR221 |
| MBAP1 ΔVK MD | TTAGCGGATCCGAAAACTCCAACTCTGCAGGA | TTTTCGGATCCGCTAAGTCTCCTTGT ATGATGAAT | pK7m34GW |
| MBAP2 CDS | GGGGACAAGTTTGTACAAAAAA GCAGGCTTAATGAGGTCAGATACGGTTC | GGGGACCACTTTGTACAAGAAAGCT GGGTTTCTTGGCTTCGAATCATTTC | pDONR221 |
| MBAP3 CDS | GGGGACAAGTTTGTACAAAAAA GCAGGCTTAATGAGACCAGCTATACCTC | GGGGACCACTTTGTACAAGAAAGCT GGGTTTTTCCCTTTGCTACGAAAATTGG | pDONR221 |
| BRI1 CDS | GGGGACAAGTTTGTACAAAAAAGCAGGCTTA ACCATGAAGACTTTTTCAAGCTTCTTTC | GGGGACCACTTTGTACAAGAAAGCT GGGTATAATTTTCCTTCAGGAAC | pDONR221 |
| BKI1 CDS | GGGGACAAGTTTGTACAAAAAA GCAGGCTTAGAAACTAATCTACAACAG | GGGGACCACTTTGTACAAGAAAGCT GGGTATCAAGAATCCTTAACCTT | pDONR221 |
| MBAP1 1-190 | GGGGACAAGTTTGTACAAAAAA GCAGGCTGGATGAAGGCTGATACTGTTCTAGAC | GGGGACCACTTTGTACAAGAAAGCT GGGTTTGGTCTTCTTTGCGATAGTG | pDONR221 |
| MBAP1 191-489 | GGGGACAAGTTTGTACAAAAAA GCAGGCTTAATGGACCTCATCACCAAGAAC | GGGGACCACTTTGTACAAGAAAGCT GGGTTTCTTGTCCTCTGTGTTTG | pDONR221 |
| MBAP1 191-489 stop | GGGGACAAGTTTGTACAAAAAA GCAGGCTTAATGGACCTCATCACCAAGAAC | GGGGACCACTTTGTACAAGAAAGCT GGGTTTTATCTTGTCCTCTGTGTTTG | pDONR221 |
| MBAP1 490-1061 | GGGGACAAGTTTGTACAAAAAA GCAGGCTTAATGCCTAGAGAAAGCCCTC | GGGGACCACTTTGTACAAGAAAGCT GGGTATCTTGGCTTCGAGTCGTTTCC | pDONR221 |
| pMBAP1 | GGGGACAAGTTTGTACAAAAAA GCAGGCTGGCCGGTGACGGAATCTTACCTCT | GGGGACCACTTTGTACAAGAAAGCT GGGTATTGGAGGGGTAAATTAGTCTCTTA | DONR-P4P1R |
| pMBAP2 | GGGGACAACTTTGTATAGAAAAGTTGTA CCAACTTGTATAAACGCCGC | GGGGACTGCTTTTTTGTACAAACTTGT TCTCACAAGACCAGATTGCTAATG | pDONR-P4P1R |
| pMBAP3 | GGGGACAACTTTGTATAGAAAAGTTGTA AGGATTTAAAGCCATGTCTCC | GGGGACTGCTTTTTTGTACAAACTTGT CTTGACCCCTTTACACAAAATTTTC | pDONR-P4P1R |

**Table S4. Final constructs used and destination vectors**

| Construct | Destination vector |
|---|---|
| 35S::MBAP1-mCherry | pK7m34GW |
| 35S::MBAP1ΔVK-mCherry | pK7m34GW |
| 35S::VK-mCherry | pG7m34GW |
| pLacI::MBP-MBAP1 | pKM596 |
| 35S::MBAP1-mCitN | pB7m34GW |
| 35S::MBAP1-mCitC | pH7m34GW |
| Ubi10::eGFP-MBAP1 | pB7m34GW |
| 35S::(1-190)MBAP1-mChe | pH7m34GW |
| 35S::(191-489)MBAP1-mChe | pH7m34GW |
| 35S::(490-1061)MBAP1-mChe | pH7m34GW |
| pLacI::MBP-MBAP1(191-489) | pKM596 |
| Ubi10::MBAP2-eGFP | pG7m34GW |
| 35S::MBAP2-mCitC | pH7m34GW |
| Ubi10::MBAP3-eGFP | pG7m34GW |
| 35S::MBAP3-mCitC | pH7m34GW |
| 35S::BRI1-mChe | pH7m34GW |
| 35S::BRI1-mCitN | pB7m34GW |
| 35S::BRI1-mCitC | pH7m34GW |
| 35S::BRL1-mCitN | pH7m34GW |
| 35S::BRL3-CitN | pH7m34GW |
| Ubi10::BKI1-mCit | pB7m34GW |
| 35S::LecRK-I.9-mCitN | pK7m34GW |
| 35S::LecRK-I.9-mCitC | pK7m34GW |
| pMBAP1::eGFP-GUS | pB7m34GW |
| pMBAP2::eGFP-GUS | pB7m34GW |
| pMBAP3::eGFP-GUS | pB7m34GW |
| pLacI::MBP | pMAL-c2X |

**Table S5. Primers used for RT-PCR**

| Gene | Forward | Reverse |
|---|---|---|
| *MBAP1* | ATTGAATCACAAGCTTCAAAGAGAG | GGCATAACAGGAGACGGTATA |
| *MBAP2* | GAGAAAAGCAACACTTGGTTCACT | ACATATTATTAGCAGCAGCAACGGT |
| *MBAP3* | AAGCAACAAAAATGCTGGAACCT | TTAAAGCTCAATGGTGTGCCAG |
| *BRI1* | TCCGCGGTGTGATCCTTCAAAT | GCCGTGTGGACCAGTTTAGTTT |
| *ACT2* | GCCCAGAAGTCTTGTTCCAG | TCATACTCGGCCTTGGAGAT |

**Table S6. Primers used for RT-qPCR experiment**

| Gene | Primer F | Primer R |
|---|---|---|
| *BAS1* | CCAAGGACCATGTCGTTAAGC | CCTGAAGTATAGCAAGATTCTGACC |
| *DWF4* | GGTGATCTCAGCCGTACATTTGGA | CCCCACGTCGAAAAACTACCACTTC |
| *CPD* | TTGCTCAACTCAAGGAAGAG | TGATGTTAGCCACTCGTAGC |
| *EF1α* | GTCGATTCTGGAAAGTCGAC | AATGTCAATGGTGATACCACGC |
| *AT1G58050* | CCATTCTACTTTTTGGCGGCT | TCAATGGTAACTGATCCACTCTGATG |
